# Genetic background and transient prenatal disruption of vitamin A signaling determine susceptibility to airway hyperresponsiveness in mice

**DOI:** 10.1101/2025.10.31.685835

**Authors:** Takehiro Otoshi, Ayyappa K.S. Kameshwar, Benjamin D. Kotton, Yoshinori Seki, Zachary Cardell, Ke Xiangyi, Yuta Matsuno, Pooja Rajaram, Youn-Kyung Kim, Sarah M. Sharpton, Loredana Quadro, Wellington V. Cardoso, Masako Suzuki

**Author notes:** Equal contribution (Wellington V. Cardoso, Masako Suzuki). **Correspondence**: Wellington V. Cardoso MD. PhD, Columbia Center for Human Development, Columbia University Irving Medical Center, Department of Medicine. 650 West 168th Street, BB 8-812, New York, NY 10032, Masako Suzuki, DVM, PhD, Texas A&M University, 498 Olson Blvd, College Station, TX, 77840.

## Abstract

Airway structural changes and hyperresponsiveness (AHR), hallmarks of asthma, are crucially influenced by genetic variations and adverse exposures. While intrauterine environmental perturbations leading to dysfunctional lung development have been linked to adult pulmonary disease, still little is known about the developmental events leading to these postnatal abnormalities. Here, we provide evidence of genetic background playing a key role in this process. Using A/J and C57BL/6J mice known for their distinct susceptibility to AHR, we show that A/J but not C57BL/6J develop an aberrant airway smooth muscle (SM) program and AHR in adulthood when exposed transiently to a vitamin A/retinoic acid (RA)-disrupted intrauterine environment *in vivo* by a maternal BMS493 administration. Single nuclei multiomics analysis identified a subpopulation of mesenchymal cells that overactivated TGFβ targets in response to BMS selectively in A/J, but not C57BL/6J, embryonic lungs. These cells, localized to sites of airway SM initiation, exhibited robust BMS-mediated upregulation of SMAD2/3 targets, including regulators of SM program *Pdgfra* and *Tnc*, and showed stable cell proportions despite the marked transcriptional rewiring following RA disruption. These findings identify TGFβ-activating mesenchymal cells as a critical niche responsive to RA signaling and reveal how genetic background determines developmental susceptibility to micronutrient perturbations with long-term impact on airway function.

## INTRODUCTION

Genetic background in combination with environmental exposures can act as a significant risk factor for a variety of human health conditions, influencing susceptibility and severity of cardiovascular, pulmonary, autoimmune disorders, and cancer (1). One of these conditions is asthma, a complex disease characterized by phenotypes that include airway hyperresponsiveness (AHR), inflammation, and structural remodeling with narrowing of the distal airways (2). AHR is clinically defined as an increased airway sensitivity to non-specific stimuli or inhaled constrictor agonists, such as methacholine (3). Although a hallmark of asthma, AHR is multifactorial and is seen as a manifestation of other pathological conditions often associated with airway structural changes that include an increase in airway smooth muscle (SM) mass. Mouse models have provided key insights into the genetic determinants of AHR susceptibility. Strain-dependent variations of AHR have been reported in different inbred mouse strains, and several AHR quantitative loci have been identified (4–9).

This AHR variation is well-illustrated from studies in C57BL/6J and A/J (hereafter B6 and AJ, respectively) mice, known for their distinct genetic backgrounds and use in pulmonary research. AJ mice are known for their high susceptibility to developing AHR and airway remodeling, while B6 mice are more susceptible to developing emphysema (10–12). Importantly, a comprehensive transcriptome analysis of the AJ and B6 lungs throughout the prenatal and postnatal stages reveals strain-related differences in gene expression signatures associated with a broad range of biological processes, even from early stages (13). This raises fundamental questions relevant to understanding the developmental origin of these abnormal airway responses. Are there differences in lung cellular behavior and composition that could ultimately reflect on distinct postnatal strain-related responses, such as AHR susceptibility and abnormal airway remodeling? How does the genetic background modulate the lung developmental programs associated with these responses?

An additional complexity to consider is the impact of maternal-fetal interactions influencing the developmental processes in a genetic background-dependent fashion. An essential component of these interactions is the establishment of an appropriate maternal micronutrient status in the intrauterine environment (14). Micronutrient perturbations have been linked to altered developmental events and postnatal adverse health outcomes (15). A large body of evidence implicates vitamin A and its active form, retinoic acid (RA), as key micronutrients regulating biological functions from early organogenesis to adulthood. In the adult lung, vitamin A/RA deficiency has been associated with airway SM dysfunction and AHR in animal models and humans (16, 17). Still, little is known about how genetic background influences these responses, likely contributing to the conflicting results on the relationship between vitamin A status and asthma risk (18). Compelling evidence of the lasting adverse effects of a brief prenatal vitamin A perturbation on adult lung function was previously reported. Mouse embryos briefly exposed to vitamin A deficient intrauterine environment develop subtle but relevant changes in airway SM structure and show AHR in adulthood (19, 20).

The distinctive ability of AJ and B6 mice to undergo airway remodeling and AHR, and the observations reported above, provided a unique opportunity to explore this model of prenatal RA perturbation to address relevant questions about the developmental origins and the impact of genetic background in this phenotype. Are the lung developmental programs similarly sensitive to perturbations in the RA status in AJ and B6? If so, what makes them different, and what cellular and molecular changes are potential determinants of these strain-specific responses?

Here, we address these questions by performing comprehensive multiomics and functional analysis of AJ and B6 mice similarly subjected to prenatal RA deficiency. By disrupting RA signaling prenatally during a short developmental window, we found remarkably different responses in lung gene expression and function between these two strains. In contrast to B6, AJ mice showed an exacerbated response to RA disruption, activating TGFβ signaling and transcriptional targets differentially in a subpopulation of mesenchymal progenitor cells. Our data show that this subpopulation is spatially distributed where the SM program emerges in the distal developing lung, and that in AJ mice, it is exquisitely sensitive to RA to maintain proper expression of TGFβ targets, such as *Pdgfra*, *Tnc,* and others required for proper SM differentiation. Collectively, these observations emphasize the effect of genetic background on the impact of developmental perturbations on adult AHR, highlighting mechanisms in the SM program that mediate such responses.

## RESULTS

### Genetic background is a determinant of the aberrant response of the embryonic lung to an intrauterine RA-deficient environment and the susceptibility to AHR in adulthood

A comprehensive analysis of the impact of genetic background on pulmonary function across 36 distinct inbred mouse strains showed remarkable differences in AHR to methacholine (MeCH) (4). AHR can result from multiple factors, including abnormal inflammatory, structural and neural responses. There is compelling evidence of an association between AHR and airway remodeling with prenatal disruption of vitamin A signaling in mice (19). We reasoned that further examining this association in mice with distinct genetic background could provide relevant insights into the developmental origins and prenatal events influencing AHR susceptibility. However, we had no evidence that the observations above, which were reported in an outbred (CD1) line, could be reproduced in mice with defined genetic backgrounds. Even less certain was the relationship between vitamin A/RA status and AHR in mice of different backgrounds. Thus, to investigate these issues we selected A/J (AJ) AJ and C57BL/6J (B6), two widely used inbred mouse strains known for their markedly distinct responses to cholinergic challenge (4). To confirm the suitability of these strains for our subsequent studies, adult 16 weeks-old male AJ and B6 mice were subjected to whole-body plethysmography (FinePoint), and specific airway resistance (sRaw) was measured under nonchallenged state (baseline) or increasing concentrations of methacholine (MeCH; 6.25, 12.5, 25 and 50 mg/ml) (21). Differences in sRaw were found in both strains in response to MeCH **(Figure 1A)**. However, AJ showed significantly higher airway resistance across MeCH concentrations compared to B6. Our findings of sRaw and AHR in AJ mice were consistent with airway resistance reported in the Mouse Phenome Database (AJ, 0.424± 0.096; B6, 0.321±0.080; p<0.05).

**Figure 1.**
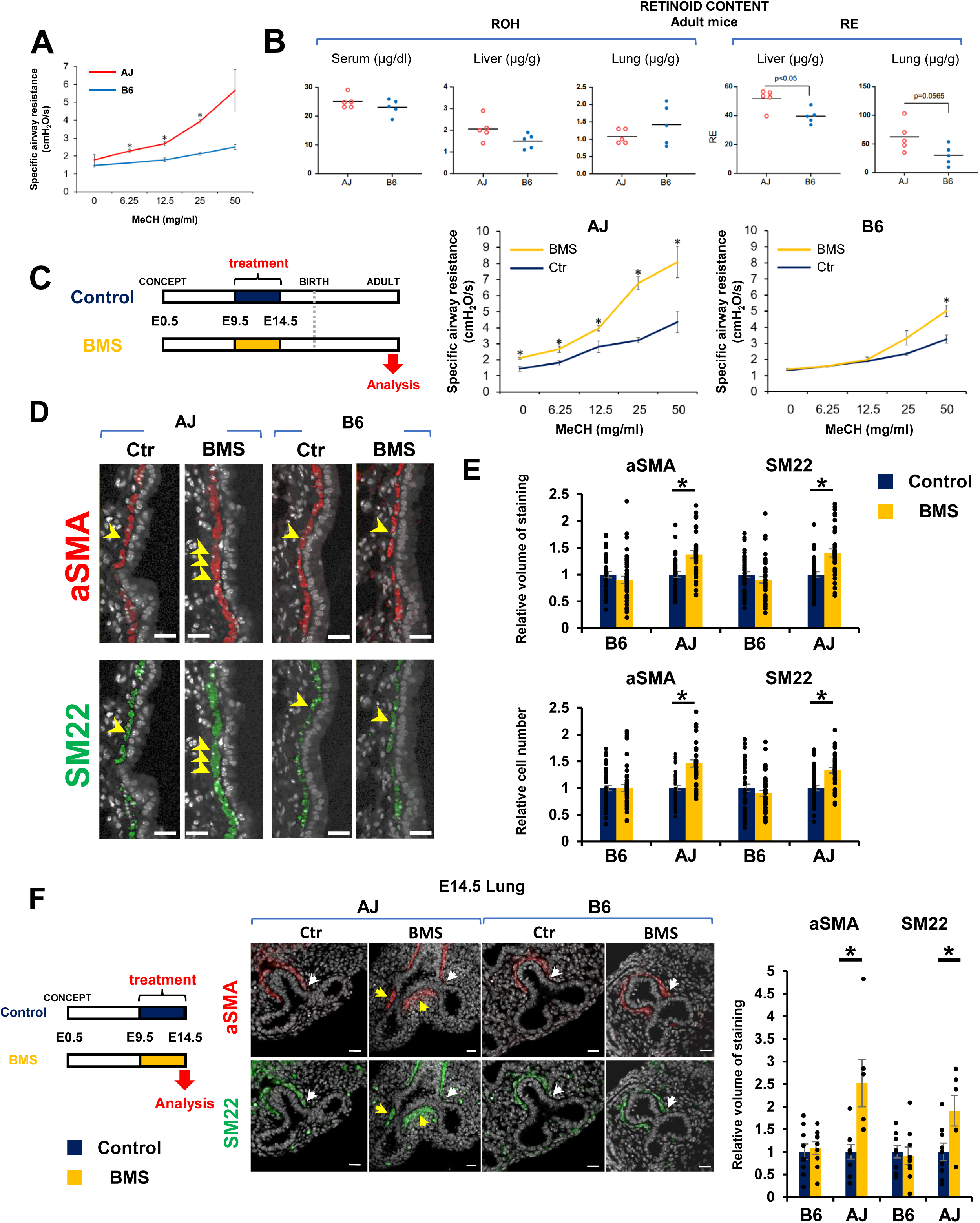
Genetic background is a key determinant of the airway smooth muscle (SM) program and airway hyperresponsiveness (AHR) in mice exposed transiently to a RA-deficient intrauterine environment. **(A)** Analysis of specific airway resistance (sRaw) in adult AJ and B6 mice under methacholine (MeCH) challenge (whole-body plethysmography; Fine Point Data Science). Significant differences in sRaw between AJ and B6 at increasing MeCH concentrations. Graph: mean ± SE (n=3 per group), \**p*<0.05. **(B)** Retinoid concentrations (retinol ROH, retinyl ester RE) in serum and tissue homogenates from lung and liver from adult AJ and B6 mice fed a control chow diet (Lab Diet, 5LG4). Graph: mean ± SE (n=5 per group). **(C)** Effect of transient prenatal disruption of RA signaling on sRaw in adult male AJ and B6 mice. Diagram experimental design (left): Maternal BMS administration (3.75 μg/body weight per day: μg/bw/day) or corn oil control from gestation days 9.5-14.5 followed by control diet onwards. Airway resistance upon MeCH challenge (right) in adult offsprings exposed prenatally to BMS or control conditions. Graphs: sRaw (mean ± SE) BMS vs control from each strain (\**p*<0.05). AJ Control (n=4), AJ BMS (n=3), B6 Control (n=9), B6 BMS (n=6). (**D and E**) Immunofluorescence (IF) and quantitative analysis of αSMA and SM22 in adult AJ and B6 lungs exposed prenatally to BMS or control conditions. Graphs: Relative volume of staining for each marker and relative number of labeled cells per basement membrane in airways in each condition. AJ Control (n=39), AJ BMS (n=37), B6 Control (n=45), B6 BMS (n=43). (**F**) Analysis of E14.5 lungs exposed to maternal BMS (3.75 μg/bw/day) or control diets. Diagram experimental procedure (left). αSMA and SM22 IF and morphometric assessment of airway SM (relative volume of staining per basement membrane) in freshly isolated lungs. Arrows: normal distribution (white) and ectopic accumulation (yellow) distribution of SM markers in AJ and B6 lungs. Graph: mean ± SE for each marker (\**p*<0.05). AJ Control (n=9), AJ BMS (n=6), B6 Control (n=9), B6 BMS (n=9). Scale bars in D and F: 20um.

Given that differences in retinol (ROH) and retinyl ester (RE) levels have been reported in tissues from different adult inbred mouse strains, and that inadequate adult vitamin A/RA status has been linked to AHR, we asked whether the distinct sRaw/AHR of AJ and B6 reflected strain-dependent differences in retinoid bioavailability (22). The concentrations of ROH and RE in serum and in homogenates from lung and liver were measured in 8-week-old adult mice under a standard VitaminA-suficient diet (Lab Diet, 5LG4). No significant differences in lung RE levels (p=0.0565) or ROH levels in lung, liver or serum were found between strains **(Figure 1B)**. Although liver RE levels were higher in AJ (p<0.05), overall, the differences in retinoid content between adult AJ and B6 mice were unlikely to explain the distinct AHR behavior of these strains.

We then tested whether AJ and B6 embryos were equally susceptible to developing AHR in adulthood when briefly exposed to a vitamin A/RA deficient intrauterine environment during early lung development (19). To ensure effective and transient disruption of RA signaling in both strains, we used a more direct approach to prevent RAR activation in the embryo by orally administrating mothers with a well-established pan-RAR reverse agonist, BMS493 (BMS) (19, 23–25). BMS administration could also minimize the variability in vitamin A uptake and retinoid bioavailability seen across strains. We conducted pilot studies to identify a BMS concentration that could simulate the effects of a mild-moderate vitamin A deficiency without the known teratogenic effects in embryonic growth or organogenesis, allowing offspring to survive to adulthood. Thus, AJ and B6 dams were daily gavaged with BMS at different concentrations (3.75-15 μg per body weight per day, μg/bw/day) or vehicle, corn oil (control group). Embryos were exposed to BMS in vivo from E9.5-14.5, a developmental window previously reported to encompass the initial stages of branching morphogenesis and airway differentiation (**Supplemental Figure 1A)**. Gross morphological analysis of E14.5 embryos exposed to the highest BMS concentration (15 μg/bw/day) showed severe effects in the body size and lung growth most evident in AJ mice. AJ lungs were severely hypoplastic compared to respective controls and B6 lungs. These abnormalities were largely undetected at 3.75 μg/bw/day. Lungs from both AJ and B6 embryos exposed to BMS at this concentration appeared macroscopically comparable to respective controls (**Supplemental Figure 1B)**.

qPCR analysis of these embryonic lungs confirmed that intrauterine exposure to BMS 3.75 μg/bw/day effectively disrupted RA signaling, as seen by the significant decrease in *Rarb and Cyp26b1* expression compared to controls (n≥3, p <0.05). Importantly, we found no downregulation in expression of the pan-epithelial marker *Epcam* between control and BMS lungs in both strains **(Supplemental Figure 1C)**. This ensured that BMS was effective in disrupting RA signaling but not lung epithelial growth and branching. To further demonstrate that the prenatal perturbation in RA signaling did not result in the teratogenic effects that prevent viability, we monitored these mice until adulthood. Embryos from both A/J and B6 mothers exposed to BMS (3.75 μg/bw/day; E9.5-14.5) were viable, comparable to controls, and reached adulthood. No difference in body weight or liver weight relative to body weight was found in mice prenatally exposed to BMS or control in both strains. Although we detected a small difference in lung weight relative to body weight between adult mice prenatally exposed to BMS compared to control conditions, this difference was similarly found in both AJ and B6 (n>3, from at least 2 dams) and, importantly, did not prevent animals to undergo postnatal development and reach adulthood as in controls **(Supplemental Figure 1E)**.

Next, we asked whether the brief disruption of RA signaling in embryos exposed to BMS differentially influenced the susceptibility of AJ or B6 mice to develop AHR in adult life. sRaw was assessed at baseline and after MeCH challenge in adult 16-week-old male mice from all groups as before. Interestingly, brief intrauterine exposure to an RA-deficient environment consistently resulted in aberrant higher sRaw at baseline and across all MeCH concentrations in adult AJ, but not B6, which responded to the challenge only at the highest MeCH concentration **(Figure 1C)**. We reasoned that the hyperresponsiveness of BMS-AJ was associated with some structural changes in the airway SM, as suspected by similar observations previously reported in CD1 mice. Indeed, immunofluorescence (IF) and quantitative analysis confirmed the increased thickness in airway SM labelled by αSMA and SM22 in the adult AJ lungs compared to B6 **(Figure 1D-E)**. To confirm the developmental origin of this phenotype, we examined embryonic lungs from both AJ and B6 mice immediately after the last day of exposure to maternal BMS administration. E14.5 BMS-AJ lungs showed no obvious abnormalities in epithelial patterning or growth; however, αSMA and SM22 IF revealed increased ectopic accumulation of airway SM, consistent with the pattern previously reported in vitamin A-deficient embryos in CD-1 mice (19). By contrast, disruption of RA signaling had no effect on SM markers in BMS lungs from B6 embryos **(Figure 1F).**

These findings were consistent with the idea that differences in genetic background are major determinants of how the lung responds to prenatal perturbations in RA signaling and the potential link to the susceptibility to AHR in adult life.

### AJ and B6 lungs differ broadly in gene expression signatures with minimal differences in cellular composition in response to prenatal RA deficiency

The distinct response of AJ and B6 embryos to prenatal disruption of RA signaling led us to examine differences in lung gene expression signatures associated with these distinct backgrounds. First, we performed bulk RNA seq in E14.5 lungs from AJ and B6 embryos intrauterine exposed to BMS or control conditions (n=4 per group). Interestingly, Principal Component Analysis (PCA) showed that the strain-related differences in gene expression between control AJ and B6 lungs were greater than those resulting from the prenatal exposure to BMS in each strain (87% variance vs. 4% variance) (**Figure 2A-B, and Supplemental Table 1**). Indeed, control AJ and B6 lungs differed greatly in their signatures, as revealed by 1258 differentially expressed genes (DEGs) (**Figure 2C, and Supplemental Figure 2B**).

**Figure 2.**
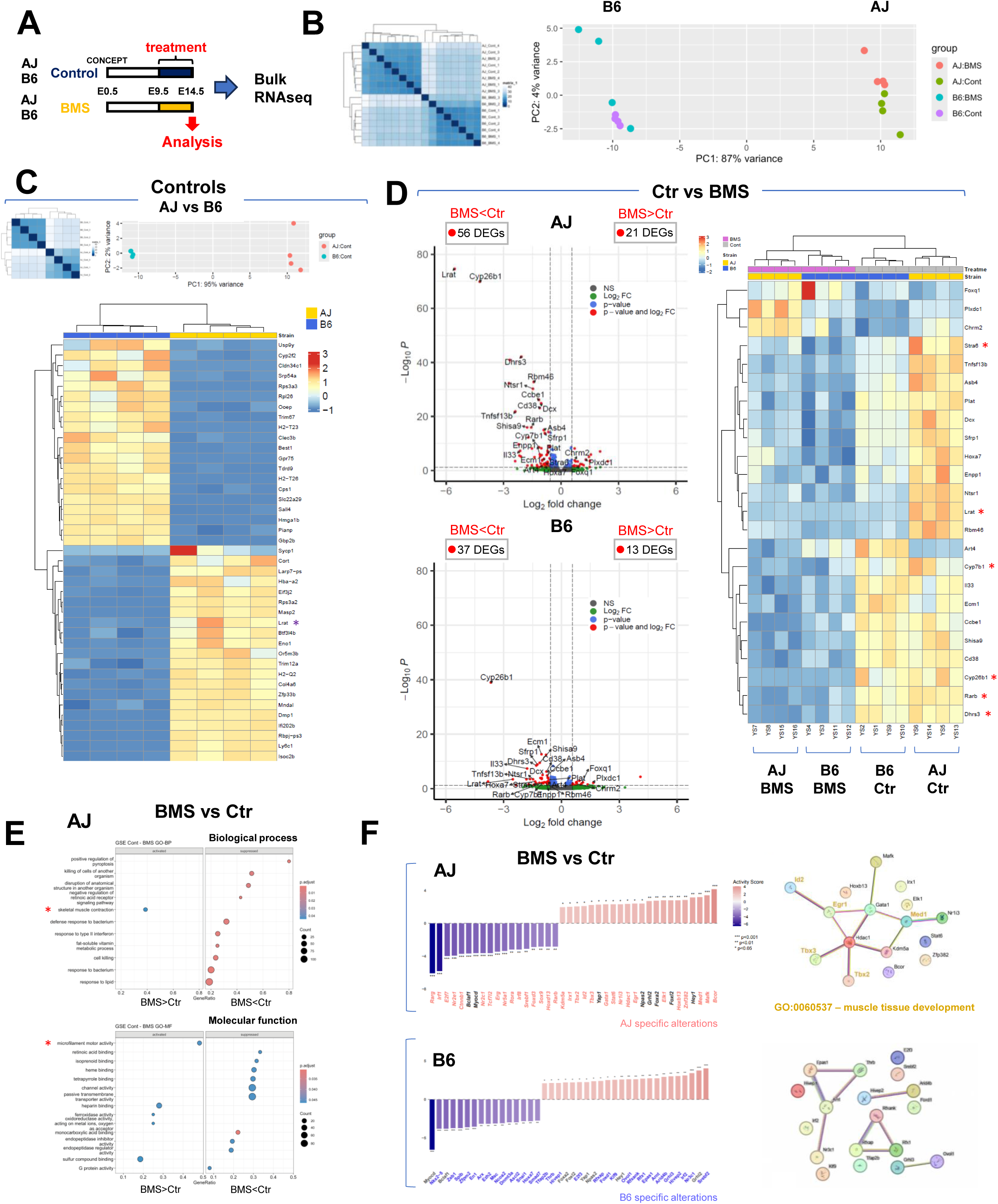
AJ and B6 developing lungs have distinct gene expression signatures and differential activation of a SM gene regulatory network under RA-deficiency. **(A)** Experimental design: E14.5 lungs from AJ or B6 embryos under maternal BMS (3.75 μg/bw/day) or control diets. **(B)** Bulk RNA-seq: Hierarchical clustering heatmap of sample-to-sample distances from variance-stabilized RNA-seq data: segregation by strain (AJ vs B6) (left). Principal Component Analysis (PCA, right) plot of variance-stabilized RNA-seq data: sample clustering based on strain and treatment status. Points colored by strain and treatment revealing transcriptional shift by BMS administration in AJ and B6 E14.5 lungs. PC1 and PC2 variance: strain effect vs BMS effect. **(C)** Differential gene expression between controls AJ and B6 E14.5 lungs. Hierarchical clustering and unsupervised heatmap depicting the top genes enriched in each strain (* depicts *Lrat* as a top enriched in AJ control, see text). **(D)** Distinct response of AJ and B6 embryonic lungs to BMS exposure. Volcano plots: differentially expressed genes (DEGs) in BMS vs control embryonic lungs from AJ and B6 mice. Number of DEGs downregulated or upregulated depicted in boxes and highlighted in red if padj < 0.05 and log2FC > 0.58. Supervised heatmap (DEGs control vs BMS from bulk RNA-seq highlighting markers of RA activity (*) downregulated in AJ and B6. **(E and F)** Gene set enrichment (GSE) and Undirected Linear Motif (ULM) analysis of transcription factor (TF) activity in BMS vs control from AJ and B6 lungs. Plots in **(E):** significantly enriched GO terms (padj< 0.05) activated (left) or repressed (right) ordered by GeneRatio in AJ lungs; (*) highlight GO terms associated with muscle function. **(F)** Analysis of TF activity: Bar charts depict the 25 most differentially activated TFs ranked by activity in BMS vs Control (AJ: top; B6: bottom). Positive scores (red): increased activity in BMS; negative scores (blue) increased activity in control (*p<0.05, **p<0.01, ***p<0.001). Right panels: String DB network visualization of key regulatory pathways altered by BMS in each strain: increased activity of muscle tissue development-related TFs in BMS-AJ but not BMS-B6 lungs. Red nodes: TFs with significantly elevated activity by BMS. Edge thickness represents confidence of functional associations based on experimental evidence, co-expression data and pathway information.

We searched for putative direct RA targets among these genes, initially by looking for RA-responsive elements (RARE)-containing open chromatin regions (RARE-OCRs) in publicly available ENCODE ATAC-seq mouse datasets from E14.5-E16.5 lungs, representative of early stages of lung differentiation (13) **(Supplemental Figure 2A,** see Methods). This revealed 28,355 RARE-OCRs with an irreproducible discovery rate <0.05, nearly half of these (44.2%) were identified within the promoter region of genes, potential RA targets (RARE-OCR genes, **Supplemental Table 2**). The data suggested a robust RA-responsive gene network at these developmental stages. The proportion of RARE-OCRs that overlapped with genes differentially expressed between control AJ and B6 lungs was not markedly different (AJ: 39.7%; B6: 54.4%). However, Gene Set Enrichment Analysis (GSEA) suggested that they differed considerably in downstream pathways and presumably function. For example, analysis of RARE-OCR-containing genes enriched in AJ included GO terms, such as *growth factor activity, lyase activity, and transferase activity,* while those enriched in B6 included *extracellular matrix-related pathways and growth factor binding* **(Supplemental Figure 2B)**. Moreover, confirming our initial findings, *Stra6* and *Lrat*, key genes associated with vitamin A uptake and retinoid bioprocessing/storage, were among the DEGs upregulated in control AJ compared to B6 **(Supplemental Figure 1C-D, and Supplemental Table 3)** (26–32).

We then compared the impact of maternal BMS exposure on gene expression signatures of AJ and B6 embryonic lungs. Efficient disruption of RA signaling was confirmed in both strains by downregulation of RA pathway target genes (*Rarb*, *Cyp26b1*, *Dhrs3*) as well as other known RARE-OCR containing genes such as *Hox* family members **(Figure 2D, Supplemental Figure 2C, and Supplemental Table 4)**. GSEA of DEGs in BMS vs control lungs from AJ embryos revealed significant enrichment for terms associated with the suppression of RA-related molecular function (ex. RA binding), as well as enrichment for terms associated with activation of muscle contraction and microfilament activity, none of these found in B6 lungs **(Figure 2E and Supplemental Figure 2D)**. Consistent with this, analysis of transcription factor activity using the R package, *decoupleR* with Undirected Linear Motif (ULM) approach showed decreased *Rarg*, *Rarb*, *Rora* activity and increased activity of transcription factors, such as *Egr1, Id2, Med1, Tbx2, and Tbx3* associated with muscle development (GO: 0060537) in AJ BMS (**Figure 2F**). This was not observed in BMS vs control B6 lungs, suggesting a less robust RA-related functional activity in these events, at least in the context of our experimental conditions. Together, the data pointed to broad differences in the gene expression signatures of the developing AJ and B6 lungs, including regulators of retinoid bioavailability and RA targets. However, these differences could hardly justify their role as determinants of the altered response of AJ mice to an RA-deficient environment.

To further investigate whether the distinct behavior of AJ and B6 resulted from predominant changes in gene expression in a specific cell population in response to BMS, we performed single nuclei multiome analysis of these lungs exposed to BMS under the same conditions. After filtering low-quality entries, 21,293 nuclei were examined across all conditions (**AJ**: Control=4,750 and BMS=6,363; **B6:** Control=5,321 and BMS=4,859). Using weighted nearest neighbors (WNN) to integrate snRNA-seq and snATAC-seq data, four populations were identified broadly as epithelial, mesenchymal, endothelial, and immune cell based on reported markers and chromatin structure profiles **(Figure 3A and Supplemental Figure 3A)** (33, 34). No significant strain-specific differences in cell proportion were found in any of these four cell populations in response to prenatal disruption of RA signaling **(Figure 3B)**. This was somewhat unexpected since the genetic background can account for subtle but often important variations in cellular composition resulting in strain-dependent differences in organ size and patterns of growth (35–37). By contrast, we found major changes in gene expression and to a lesser extent open chromatin accessibility in response to BMS in both strains. Consistent with our bulk RNA-seq findings, BMS exposure resulted in a higher number of DEGs in AJ lungs (n=548) compared to B6 (n=214). Moreover, BMS-AJ also had a higher number of differentially accessible regions (DARs) (n=117) than BMS-B6 (n=71); of these, 61/117 (52%) and 37/71 (52%), respectively, harbored RARE-OCRs **(Supplemental Figure 3B and Supplemental Table 5-6).** The lower number of DARs compared to DEGs, suggested that differences in transcription factor expression were more likely to be the primary driver of the RA-mediated events rather than shifts in the epigenetic state. An initial analysis of the top DEGs in each of these populations was unremarkable. The collectively higher number of DEGs and DARs in the mesenchymal cell population of AJ compared to B6 and the AJ distinctive SM differentiation program in response to BMS, led us to further focus our analysis on the lung mesenchyme.

**Figure 3.**
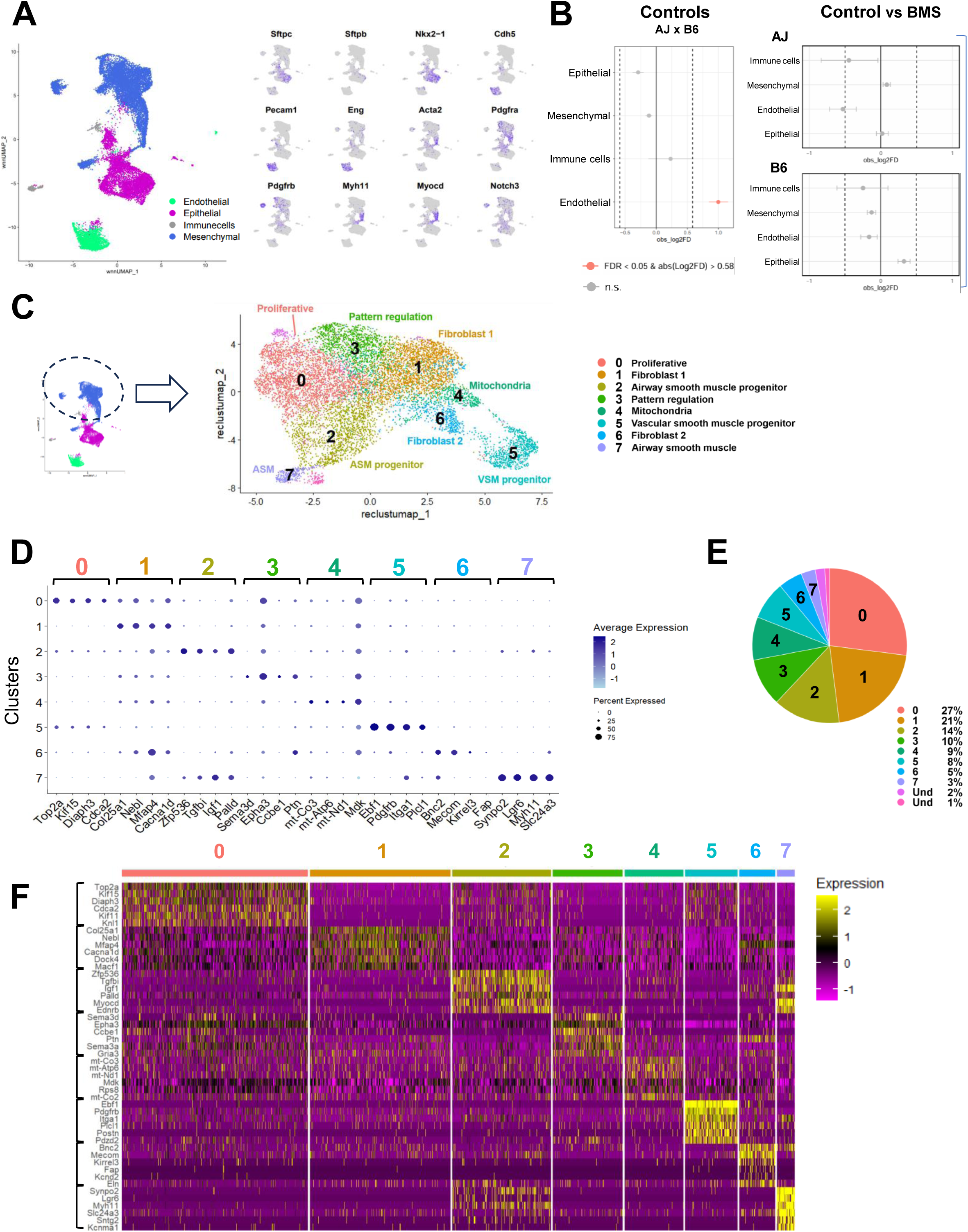
Single-nuclei multiomics shows similar shifts in cell proportion in BMS-exposed AJ and B6 and identifies multiple subpopulations of mesenchymal cells during lung morphogenesis. **(A)** UMAP visualization of weighted nearest neighbor (WNN)-integrated single-nucleus multiome data from AJ and B6 E14.5 lungs showing the main cell populations (mesenchymal, epithelial, endothelial and immune cells) identified by established markers (right panel). **(B)** Relative cell type proportions in AJ and B6. No significant differences in proportion of epithelial, mesenchymal, endothelial or immune cell populations in response to BMS (permutation testing n = 10,000). Significant differences defined by cutoffs of FDR; 0.05 and log2fold difference > 0.58. **(C)** UMAP projection of re-clustered mesenchymal cells from single-nucleus multiome analysis of E14.5 lungs (all conditions and strains). Eight subpopulations identified by established markers (FindMarkers, Wilcoxon Rank Sum) and two additional unassigned small clusters of ambiguous fate not further analyzed. **(D, E, F)** Dot plot and heatmap of top differentially enriched genes for each of the mesenchymal cluster described above. Pie chart (E) depicting the relative proportion of each mesenchymal cell subpopulation.

### Distinct subpopulations emerge from the mesenchymal cell compartment during airway morphogenesis in both AJ and B6 lungs

Given the higher number of DEGs and DARs in the mesenchymal compartment of BMS-AJ compared to BMS-B6 **(Supplemental Figure 3B),** the known diversity of lung mesenchymal cell types and the dynamic morphogenetic changes taking place in the E14.5 lung, we performed a comprehensive analysis of this compartment to understand how BMS influenced its gene expression and behavior. First, we computationally filtered for the mesenchymal cells from our multiome datasets, combining all groups. Leiden clustering identified eight subpopulations, annotated based on established markers (Seurat *FindMarkers with* Wilcoxon Rank Sum test) and two other small indistinct subpopulations with mixed signatures, not further analyzed **(Figure 3 C-F, and Supplemental Table 7).** *<u>Cluster 0</u>* was identified as undifferentiated proliferative cells for their enrichment in cell-cycle related genes (*Top2a*, *Kif15*, and *Mki67)* **(Supplemental Figure 4A)***; <u>Clusters 1</u>* <u>and *6*</u> consisted of two populations of fibroblasts-like cells based on their expression of extracellular matrix-related genes (Fibroblast 1: *Col25a1*, *Mfap4* and Fibroblast 2: *Bnc2*, *Fap*) **(Supplemental Figure 4C)**, *<u>Cluster 4</u>* was annotated as ‘mitochondria-enriched mesenchyme’ (*mt-Co3*, *mt-Atp6* and *mt-Nd1)* **(Supplemental Figure 4B)***. <u>Cluster 3</u>* was characterized by enrichment in genes associated with biological processes, such as cell migration, morphogenetic movements and chemoattraction reported mostly in neuronal studies (*Sema3d*, *Epha3*, *Ptn*), but also featuring key regulators of lung pattern formation and mesenchymal differentiation (*Wnt2, Fgf10, Rspo2, Ctnna2*) **(Supplemental Figure 5A)**. Two populations *(<u>Clusters 5 and 7)</u>* have common expression statuses of genes associated with the SM program (**Supplemental Figure 4E)**. However, <u>Cluster 5</u> was annotated as ‘vascular smooth muscle’ (pericytes) based on the differential enrichment in *Pdgfrb*, *Heyl*, and *Notch3*, compared to <u>Cluster 7</u> ‘airway smooth muscle’ identified by markers, such as *Lgr6, Actg2*, *Cdh4, Acta2, Myom1, Chrm2* and others **(Figure 4B and Supplemental Figure 4 D-E)**. Lastly, *<u>Cluster</u> <u>2</u>* was enigmatic as it encompassed a population of undifferentiated mesenchymal progenitors marked *by Zfp536, Tgfbi, Igf1,* but also expressing early stages of the smooth muscle cell program (*Myocd* and *Myh11)*, overall distinct from clusters 5 and 7 **(Figure 4A)**.

**Figure 4.**
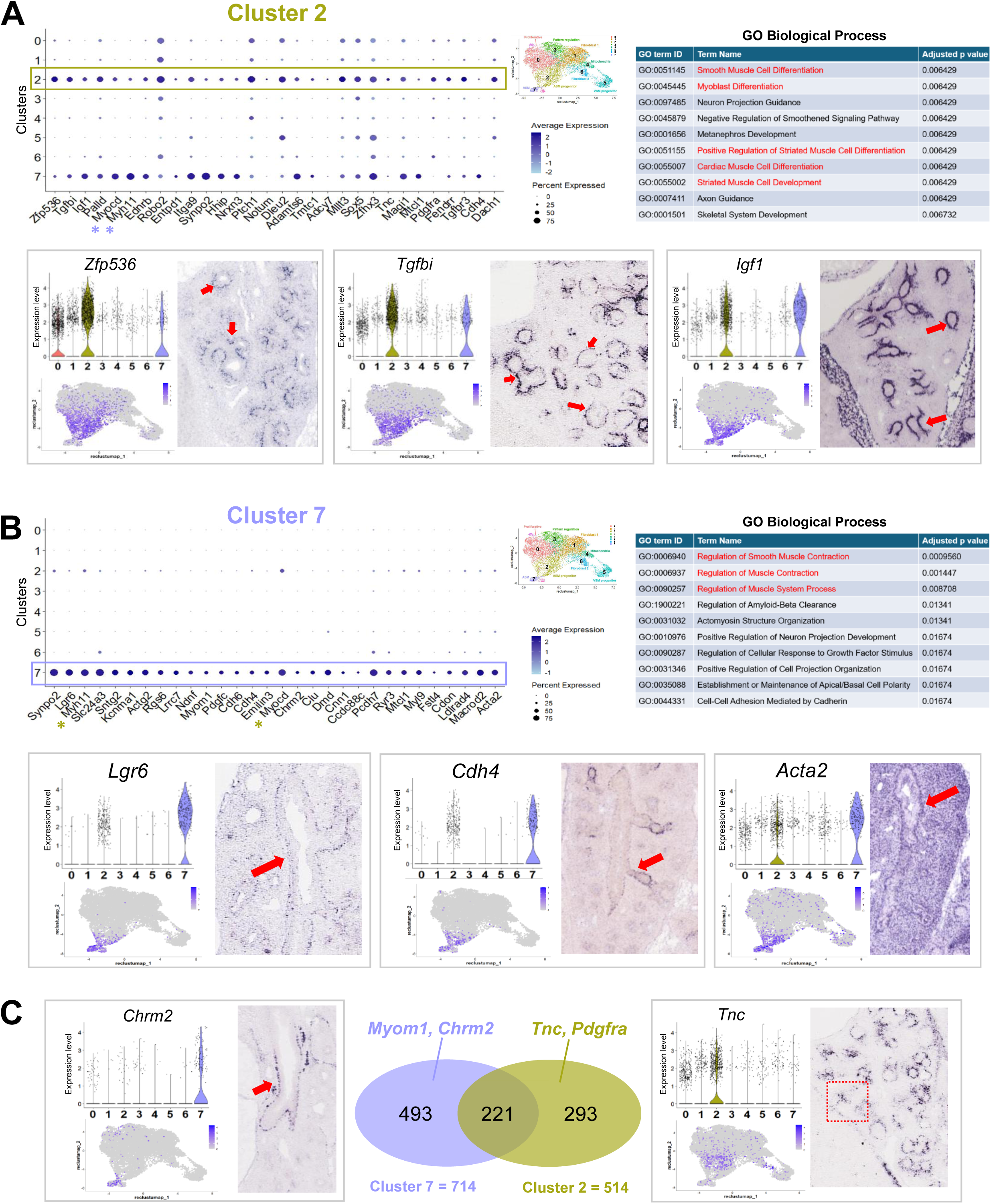
Multiomiocs analysis identifies two distinct stages of airway SM cell program during lung morphogenesis. **(A and B)** Cluster 2 and Cluster 7: dot plots displaying top DEGs (adj p-value <0.05) and Gene ontology analysis of their signatures. Enrichment in GO terms related to muscle cell development (cluster 2) and differentiation towards a contractile phenotype (cluster 7) highlighted in red. Representative genes from each cluster are depicted in violin and feature plots. In situ hybridization (GenePaint.org): distinct spatial distribution of these populations in distal mesenchyme where the SM emerges (cluster 2) or in already established airway SM at E14.5. **(C)** Venn diagram and violin plots depicting unique and shared genes between clusters 2 and 7. *Chrm2 and Tnc* are highlighted to show distal and proximal distribution (indicated by arrow and dashed box).

A systematic survey of the spatial localization of top markers from each cluster showed partially overlapping but also distinct expression patterns in the E14.5 developing lung. Transcripts from **Cluster 0** markers were found throughout the lung mesenchymal compartment, consistent with the broad distribution of proliferative cells at this stage, as also revealed by feature plots and violin plots **(Supplemental Figure 4A)**. This contrasted with the more spatially restricted distribution of cluster 2, 3, 5, and 7 genes **(Figure 4A-B, Supplemental Figure 4D-E, Supplemental Figure 5A and 5C**). Notably, analysis of **Cluster 2** markers revealed a remarkable pattern of transcript distribution in the mesenchyme at the stalk region of distal buds from where the smooth muscle program emerges **(Figure 4A).** Based on its signature, gene ontology, and spatial distribution, **cluster 2** was annotated as ‘airway smooth muscle progenitor’, reflecting its enrichment in genes associated with initiation of the cell fate program toward this lineage. Indeed, evidence from multiple studies, as well as our GSEA/GO results, showed **Cluster 2** top differentially enriched genes, components of the TGFβ, IGF, and PDGF pathways (*Tgfbi*, *Pdgfra, Igf1)* reported as early regulators of the smooth muscle program (38–41). Also enriched in Cluster 2 we found *Ptch1* and *Hhip* components of the Shh pathway reported as crucial for initiation of the airway SM program (42). By contrast, **Cluster 7** cells were most distinctly identified by the expression of more definitive markers of SM differentiation and for their localization to proximal airways (**Figure 4B-C).** The analysis also underscored the closer relationship in the signature of **cluster 7 and 2** compared to the others. For example, among the 714 and the 514 genes identified as markers of cluster 7 and 2, respectively, 30% or more genes were common to both clusters (221 genes; p-adj <0.05, absolute log2 fold > 0.585) **(Figure 4C, and Supplemental Table 14)**. These overall (Ex. *Myocd*, *Myh11)* differed in levels or percent of cells expressed in clusters 7 or 2 and likely include cells undergoing a transition from commitment to initiation of the SM program in the highly dynamic niche of the E14.5 distal lung (**dot plots in Figure 4A and 4B**). By contrast, clusters 7 and 5, in spite of being identified as SM (airway and vascular, respectively), shared only ∼10% of their DEG signatures **(Supplemental Figure 4E)** (43). Notably, DEGs in **Cluster 3** overall showed a characteristic spatial transcript distribution in the distal mesenchyme, often in a subpleural location and found collectively by GO to be regulators of cell migration, morphogenetic movements, and chemoattraction, widely reported in neurons and the lung. Additional **Cluster 3** enriched genes, such as *Wnt2 and Fgf10*, are known for their prominent role in lung pattern formation and shown by lineage tracing analysis to label early progenitors of the airway SM program **(Supplemental Figure 5A and 5B**) (44). Moreover, Wnt2 has been shown to regulate the airway SM program through MRTFB/Myocadin and FGF10 (45).

Altogether, from these annotations we propose a developmental model that aligns well with that proposed by Kumar et al. (44), which describes a three-stage progression from undifferentiated tip bud mesenchymal progenitors (Cluster 3) to recruitment to the airway SM lineage around the stalk region of distal lung buds (cluster 2) to ultimately differentiate into mature smooth muscle cells (cluster 7) **(Supplemental Figure 5D).**

### Endogenous RA signaling is differentially required in a specific TGFβ activating mesenchymal progenitor population to prevent aberrant SM program of AJ lungs during morphogenesis

Having identified the signature characteristic of these mesenchymal clusters and their spatial distribution in the E14.5 lung, we then asked what differences in cellular composition and gene expression could be attributed solely to the genetic background of each strain, selectively in the mesenchymal cell compartment. For this, we compared the gene expression signature of E14.5 AJ and B6 under unperturbed (control) conditions. Transcriptome analysis showed that AJ and B6 control lungs differed significantly in the number of DEGs in cluster 0 (478), cluster 2 (273), and cluster 3 (161), with fewer differences in the remaining mesenchymal clusters **(Supplemental Figure 6A and Supplemental Table 10).** Although interesting, we could not rule out that this could be in part due to an overall higher number of cells in these clusters. Notably, we had no evidence that endogenous RA signaling was preferentially more active in the mesenchymal populations of control AJ compared to B6 lungs. *Rarb,* a surrogate marker of RA activation, was similarly expressed in all clusters of controls from both strains **(Supplemental Figure 6B).** A comparison of the gene expression signature of clusters 0, 2, and 3 from control AJ and B6 lungs revealed *Mgp, Negr1, Col1a1*, *Opcml*, among the top DEG significantly downregulated in AJ in these three clusters. This signature was also found in AJ controls when all mesenchymal clusters were examined collectively (clusters 0-9) and when AJ and B6 were analyzed by bulk RNA-seq of whole lung homogenates. These results strongly suggest that these genes represent key components of the signatures that distinguish AJ from B6 in E14.5 lungs, ultimately associated with the strain differences in genetic background **(Supplemental Figure 6C-F)**.

Next, we assessed the impact of intrauterine exposure to BMS on the gene expression signature of AJ and B6 developing lungs. Single nuclei multiomics showed a significantly higher impact of BMS on the transcriptomics of AJ lungs. Altogether, the number of DEGs between control and BMS in all clusters was almost twice as high in AJ compared to B6 lungs (n = 142 and n = 78, respectively), the majority of these found in clusters 0, 2, and 3, regardless of differences in cell proportions **(Figure 5A and B, and Supplemental Table 8 and 9).** We leveraged the statistical power of a global analysis collectively of all mesenchymal clusters from our AJ and B6 single nuclei transcriptome dataset to identify DEGs between control and BMS-exposed lungs. Although mesenchymal disruption of RA signaling was confirmed by downregulation of *Rarb* in both strains, strain-specific differences in gene expression in response to BMS were clearly detected between AJ and B6. These included downregulation of additional components of the RA pathway (*Lrat* and *Dhrs3*), accompanied by upregulation of TGFβ targets and known regulators and markers of SM program in the mesenchymal population of BMS-AJ lungs **(Figure 5C and Supplemental Table 11).**

**Figure 5.**
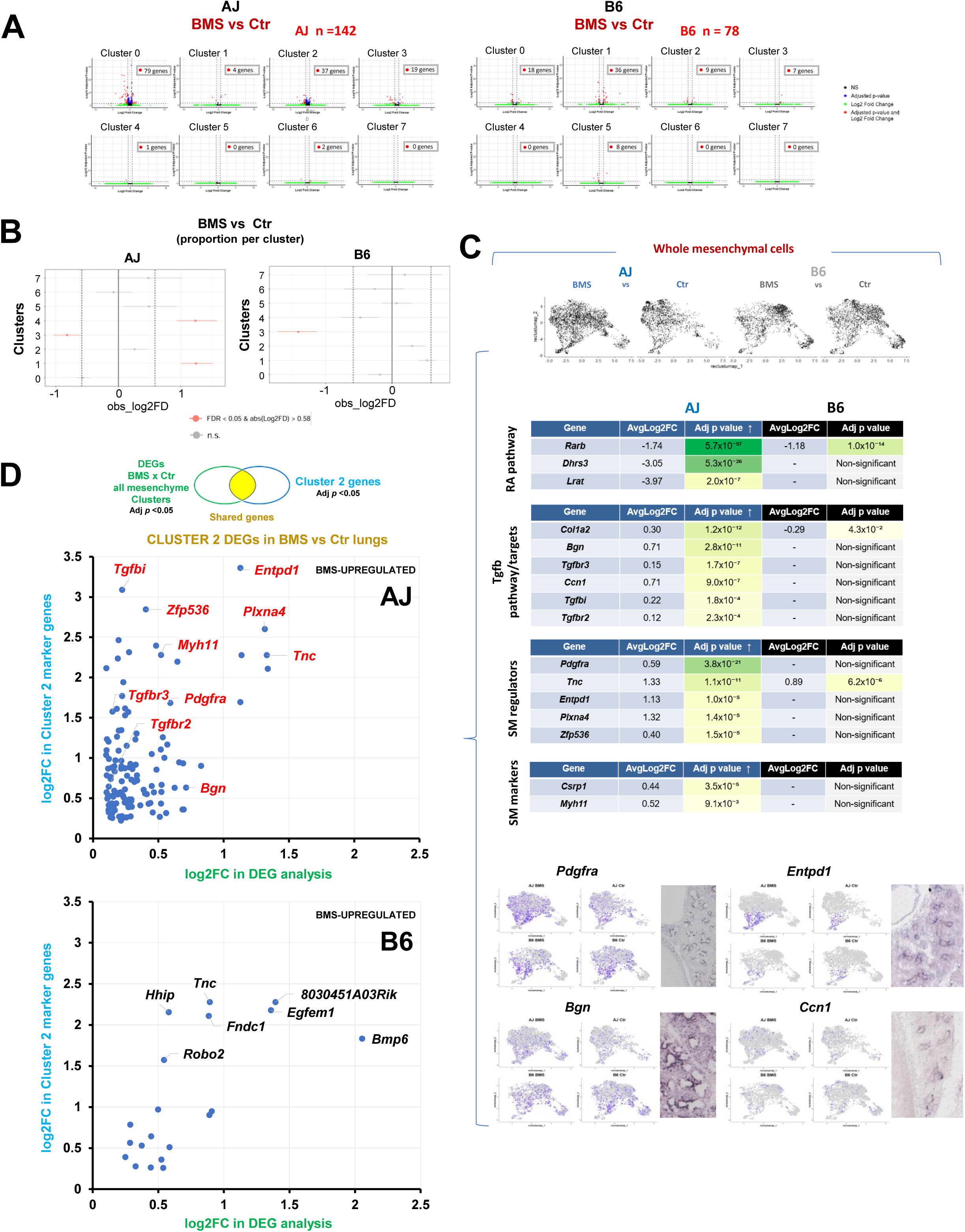
Cluster 2 cells are enriched in regulators of SM program and respond differentially to an RA-deficient intrauterine environment in AJ and B6 lungs. **(A)** Single nuclei multiomics analysis of BMS vs control in AJ and B6 lungs: Volcano plots: Higher number of DEGs in AJ, particularly in clusters 0, 2 and 3. Boxes depict DEG number with |log₂(fold change)| >0.58 and adjusted *p*-value < 0.05 relative to control in each cluster. **(B)** Graph depicting changes in cell proportion in response to BMS for each mesenchymal cluster from the E14.5 lungs in each strain. Differences between AJ and B6 are significant if FDR < 0.05, log_2_fold > 0.58, as determined by permutation testing (n = 10,000). **(C)** Analysis of the lung mesenchymal compartment (all mesenchymal clusters) identifies major differences in gene expression between AJ and B6 in response to prenatal disruption of RA signaling. Top: Diagrammatic U-map representation and tables of DEGs (BMS vs control) confirming disruption of RA signaling in both strains (*Rarb*) but distinct upregulation of markers and regulators of the SM differentiation program in AJ, compared to B6. Bottom panels: U-Map projections and spatial distribution of representative DEGs (table) showing expression associated with emerging SM program. **(D)** Identification of mesenchymal cluster 2-enriched genes among the DEGs upregulated by BMS in AJ lungs. Graphs represent the log₂(fold change) in expression between BMS and control conditions across all mesenchymal clusters from whole mesenchymal analysis (x-axis) against the log₂(fold change) of genes selectively enriched in cluster 2 (y-axis).

Given our goal to identify potential regulators of the strain-specific responses of the mesenchymal cells to an RA-deficient environment, we established a series of more stringent criteria for further analyses: a) we examined the DEGs unique to AJ not differentially altered by BMS in B6 lungs, b) among these, we focused on the genes upregulated by BMS in AJ consistent with our original hypothesis that endogenous RA prevents an unrestrained aberrant program of SM progenitors during lung morphogenesis, and, c) using information gathered from our cluster analysis and expression pattern, we selected DEGs whose mRNA distribution mapped to putative sites of initiation of the SM program in the E14.5 distal lungs. Based on these criteria, Cluster 2 emerged as the cell population to be the prime candidate for analysis. We first asked which of the DEGs identified by analysis of control vs BMS collectively in all mesenchymal cells featured among the markers of Cluster 2 cells. We then identified the cluster 2 genes differentially upregulated by BMS in AJ, but not in B6. This revealed *Tgfbi, Pdgfra, Entpd1, Zfp536* and other top markers of cluster *2* **(Figure 5D)**. The identification of widely reported regulators of the SM program among these genes further supported the criteria for further screening (38–40, 46, 47). Since BMS did not alter significantly the proportion of cluster 2 cells in AJ or B6 lungs **(Figure 5B),** we concluded that the upregulation of the cluster 2 genes found in AJ likely resulted from their de-repression in response to the disruption of RA signaling selectively in this strain. Cluster 2 genes downregulated by BMS are shown in **Supplemental Figure 7B**.

A large number of studies confirm the essential role of TGFβ signaling in SM development and in remodeling of the injured lung in diseases such as pulmonary fibrosis and asthma (39, 48, 49). The regulation of TGFβ signaling by RA in airway SM differentiation has also been extensively reported both in the developing and the adult lung. Disruption of RA signaling leads to hyperactivity of the TGFβ pathway at the onset of lung development and later during branching morphogenesis (23, 50). Here, we found key components of the TGFβ pathway enriched in the mesenchymal compartment of the E14.5 lungs **(Figure 6A).** Particularly, *Tgfbi and Tgfbr3* were top markers of cluster 2. *Tgfbr3* is a co-receptor crucial for activation of TGFβ -SMAD signaling through TGFB2-TGFBR2 binding and phosphorylation of TGFBR1 (51). There is accumulated evidence that *Tgfbi* identifies sites of canonical TGFβ activation (52, 53). First, we found strong phosphorylated SMAD2-3 signals in the stalk mesenchyme of E14.5 lungs where airway progenitors (*Sox2*+) emerge and the SM program initiates. Second, *Tgfbi* expression was markedly downregulated in lung embryonic explant cultures when endogenous TGFβ signaling was inhibited by SB431542 (SB4) **(Figure 6B).** Given the paramount importance of TGFβ signaling in regulating mesenchymal programs and evidence of the differential upregulation of TGFβ pathway genes by BMS in AJ lungs, we raised the possibility that *Tgfbi*+ cells may harbor key mediators of this differential response.

**Figure 6.**
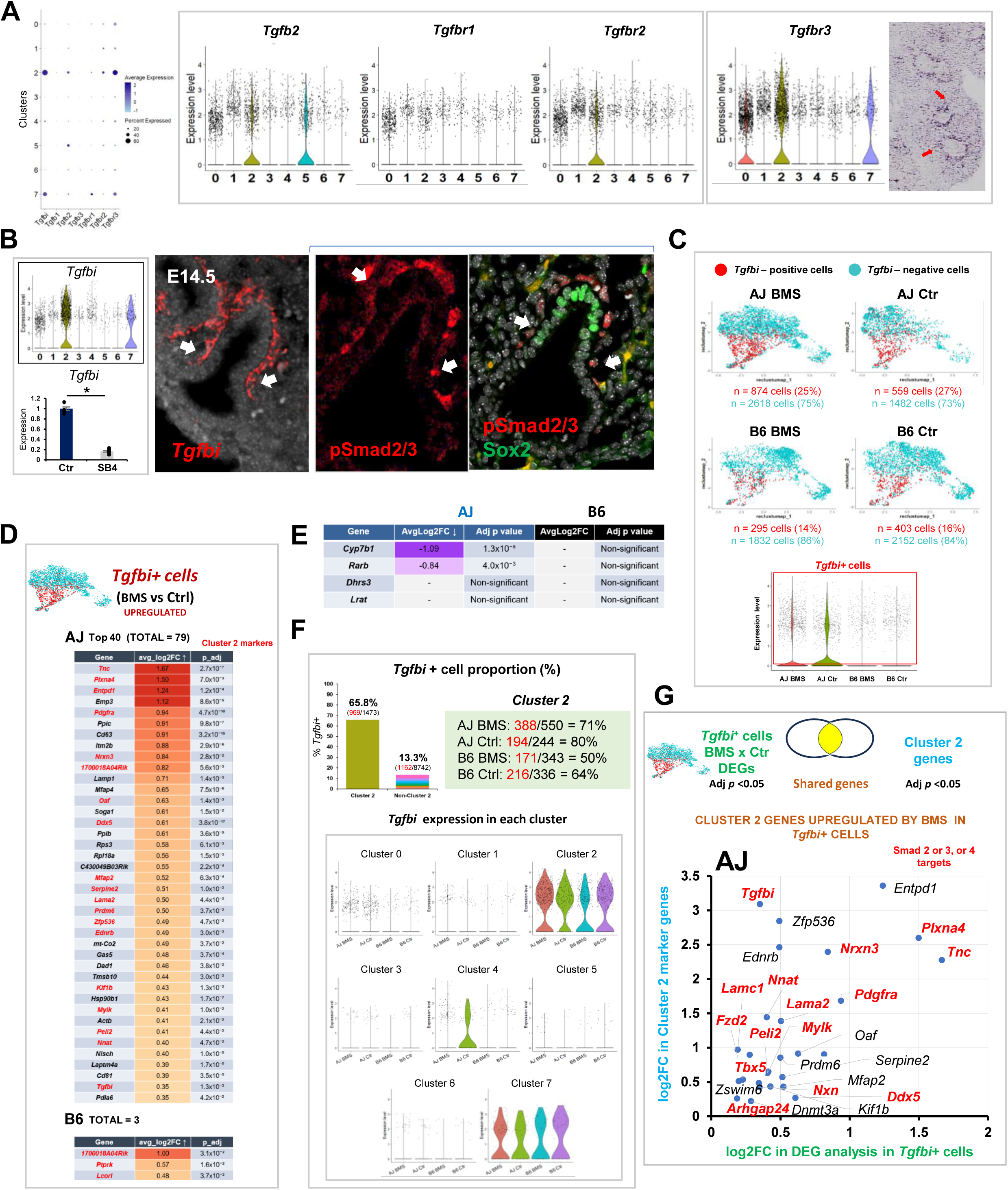
Tgfb activating cells are key determinants of the distinct response of AJ and B6 to prenatal disruption of RA signaling. **(A)** Tgfb pathway components expression across mesenchymal clusters. *Tgfb2*, *Tgfbr2*, and *Tgfbr3* enrichment in cluster 2 visualized by dot plots and violin plots. In situ hybridization (ISH): *Tgfbr3 signals* at prospective sites of airway SM initiation in distal mesenchyme (arrows). **(B)** *Tgfbi* identifies sites of Tgfb signaling activation in distal mesenchymal progenitors during lung morphogenesis. Violin plot: *Tgfbi* enrichment in clusters 2 and 7. Graph: *Tgfbi* expression in lung explants cultured under control or SB431542 (SB4)-containing media for 48h. Marked downregulation by disruption of endogenous Tgfb signaling. qPCR: mean ± SE, Ctr (n=7), SB4 (n=8). Panels: left: *Tgfbi* transcript distribution in distal E14.5 mesenchyme (RNAscope), and phosphorylated Smad2-3 IF signals in the stalk mesenchyme associated with the emergence of Sox2+ airway epithelial progenitors, where *Tgfbi* and other cluster 2 components are expressed, and the SM program initiates (see also Figure 4A). **(C)** UMAP representation of the *Tgfbi*+ cell population in AJ and B6 control and BMS-exposed lungs. The *Tgfbi*+ cell proportion differs between AJ and B6 controls, but remains largely unchanged by BMS in both strains (FDR <; 0.05 in permutation testing, n=10,000). Violin Plot: *Tgfbi* expression in mesenchymal cells from all groups. **(D)** Differential gene expression analysis of BMS vs Control *Tgfbi*+ cells from AJ and B6 lungs. Table depicts top 40 of the 79 differentially upregulated genes in AJ contrasting with only 3 genes upregulated in B6. Cluster 2 genes are highlighted in red. **(E)** Markers of RA activation (Rarb and Cyp7b1) in *Tgfbi*+ cells showing downregulation by BMS in AJ but not in B6 (padj <0.05). **(F)** Quantitative analysis of the *Tgfbi*+ cell proportion in cluster 2 compared to all other clusters and effect of BMS. Highest *Tgfbi* expression in cluster 2 and 7. **(G)** Graph depicting cluster 2-enriched genes among the DEGs upregulated by BMS in *Tgfbi*+ cells of AJ lungs (adjusted p-value <;0.05 and log2 Fold Change > 0). Differences in gene expression in BMS vs control *Tgfbi*+ cells (log₂fold change; x-axis) against the log₂(fold change) of genes selectively enriched in cluster 2 (y-axis). Genes carrying Smad2, Smad3, or Smad4 binding sites (marked in red) found upregulated by BMS in *Tgfbi*+ cells from AJ lungs.

Although largely represented in the cluster 2 population, *Tgfbi* was also found to a lesser extent in other clusters (66% of 1473 cluster 2 cells, 13% of 8742 non-cluster 2 cells) **(Figure 6B, C, F).** Thus, we computationally filtered all *Tgfbi*+ cells and collectively investigated their gene expression signature in each experimental condition. *Tgfbi*+ cells were filtered by selecting cells with normalized *Tgfbi* expression > 0.1. Analysis of control lungs from both strains showed a relatively higher proportion of *Tgfbi*+ cells in AJ (27%) compared to B6 (16%). This proportion was largely unchanged by BMS when compared to controls from respective strain (AJ BMS: 25%; B6 BMS: 14%). By contrast, the number of DEGs in BMS vs control *Tgfbi*+ cells was remarkably different in AJ (n= 337), compared to B6 lungs (n=6) (adjusted p-value <0.05) **(Supplemental Figure 7A, and Supplemental Table 13)**. Notably, while we found 79 DEGs upregulated by BMS in *Tgfbi*+ cells from AJ, only 3 DEGs were upregulated in BMS-B6 lungs **(Figure 6D)**. Given that the proportion of *Tgfbi*+ cells was unchanged by BMS in both strains, we conclude that disruption of RA signaling resulted in overly activated TGFβ signaling selectively in this cell population of AJ lungs. In searching for a potential reason for this discrepancy, we examined a panel of markers of RA activation in *Tgfbi*+ cells from both strains. Surprisingly, although *Rarb and Cyp7b1* were consistently expressed in *Tgfbi*+ cells from controls in both AJ and B6 lungs, these genes were downregulated by BMS only in AJ lungs **(Figure 6E)**. This suggested that the *Tgfbi*+ cell population of B6 was selectively insensitive to BMS and maintained its normal program, while the same population in AJ lungs under the same conditions responded aberrantly derepressing genes whose levels would be normally controlled by endogenous RA in these cells. Their identification as TGFβ activating cells in both strains and their distinct response to an RA-deficient environment are consistent with the idea that the *Tgfbi*+ cells represented the key cellular target of the SM phenotype observed in BMS AJ lungs.

To further refine our analysis and enrich genes with a greater chance of being mediators of the BMS aberrant phenotype, we first identified DEGs from cluster 2 that were most upregulated by BMS in AJ *Tgfbi*+ cells (adjusted p-value <0.05 and log2 Fold Change > 0). This ensured a greater representation in genes predicted by GSEA/GO analysis of cluster 2 to be regulators of the SM program. Second, given that *Tgfbi*+ cells were identified for their differential activation of TGFβ signaling, we used the public dataset (ChEA Transcription Factor Targets 2022) (54) to search for targets of SMAD2, SMAD3, or SMAD4 among these genes. As predicted, *Tgfbi* featured as one of the top DEGs in this cell population, further supporting this gene as a direct target of TGFβ signaling in these cells. Top differentially upregulated SMAD2-3 targets in *Tgfbi*+ AJ cells included *Tnc*, *Pdgfra*, *Plxna4*, *Nrxn3*, and *Nnat* **(Figure 6G)**. At least two of these targets, *Tnc* and *Pdgfra* are specifically relevant in the context of the question addressed here, as they have been extensively reported as part of a regulatory network implicated in SM differentiation, which encompasses interactions with the RA and TGFβ pathways. During lung morphogenesis, *Pdgfra+* cells are found abundantly in the mesenchymal compartment, including the domain of putative SM progenitors we identified in cluster 2. Loss of function studies show *Pdgfra* is essential for alveolar septation (55). *Pdgfra*+ cells in both peribronchiolar and alveolar regions differentiate into contractile cells expressing aSMA during embryonic development (40, 56, 57). Pdgfra has been shown to be elevated in the lungs of asthmatic subjects, contributing to airway remodeling and SM proliferation (58, 59). There are several lines of evidence supporting interactions of *Pdgf*, *Tgfb*, *Tnc* in myofibroblast/fibroblast differentiation in different tissue contexts. For example, a functional TGFβ-PDGFRA signaling cross-talk regulates myofibroblast migration and differentiation (60). Interestingly, a mechanism in which PDGF and TGFβ signaling promotes TNC expression in intestinal myofibroblasts has been described in a mouse model of intestinal injury (61). This crosstalk also involved SMAD3 and SMAD4, which were shown to activate the TNC promoter activity, and their overexpression further enhanced the effect of TGFβ (62). In addition, we found *Plxn4* among the most upregulated genes in *Tgfbi*+ cells. This gene encodes a receptor for semaphorins, known regulators of migration and neuron guidance. *Plxn4* is a TGFβ-SMAD3 target upregulated in SMC and fibroblasts by *Tgfb1* (63, 64). The finding of this gene enriched in this population of cells of AJ BMS lungs is intriguing since PLXN4 has been reported to be upregulated in a cohort of patients with refractory asthma (65), as well as being found to be a locus associated with low FEV1 in asthmatic patients by GWAS (66). None of the cluster 2 genes upregulated by BMS in *Tgfbi*+ cells in B6 carried Smad2, Smad3, or Smad4 binding sites (**Supplemental Figure 7C**).

Our data identifies a mesenchymal cell niche exquisitely sensitive to RA during initiation of the SM program in the developing lung of AJ mice. We show that differences in genetic background profoundly influence this mesenchymal niche, as seen by the distinct strain response to RA disruption. Collectively, these observations emphasize the effect of genetic background on the impact of developmental perturbations on adult AHR, highlighting mechanisms in the SM program that mediate such responses.

## DISCUSSION

Here, we investigated the impact of genetic background on the ability of the embryonic lung to undergo a normal program of airway SM differentiation in the presence of a transient RA-deficient intrauterine environment, and the consequences of this prenatal micronutrient perturbation on postnatal lung function and its association with AHR in adult life. Using two well-characterized inbred mouse strains, AJ and B6, with contrasting susceptibility to develop AHR, a hallmark of asthma in humans, we demonstrate that inhibiting RA signaling during an early developmental window that encompasses initiation of the airway SM program results in transcriptional rewiring of a selected population of *Tgfbi⁺* mesenchymal progenitors in AJ lungs. This cell population resides at the stalk mesenchyme of distal lung buds, where epithelial progenitors transition from SOX9 to SOX2 expression and the airway SM program is first engaged. Despite similar proportions of this TGFβ activating population in both strains, disruption of RA signaling in AJ results in robust upregulation of downstream SMAD2/3 transcriptional targets, including *Pdgfra* and *Tnc,* and in the acquisition of an aberrant differentiation signature in these SM progenitors. These changes are not seen in the B6 embryonic lungs similarly exposed to an RA-deficient intrauterine environment. The observations provide evidence of a previously unsuspected mechanism of how the genetic background influences the transcriptional landscape of the developing lung and its sensitivity to environmental perturbation.

RA and TGFβ pathways intersect extensively during lung morphogenesis (23, 50). A central mechanistic insight emerging from our work is the AJ-specific hyperactivation of TGFβ signaling in *Tgfbi*⁺ progenitors following RA inhibition. In B6, these same cells maintained expression of RA targets and showed minimal transcriptional response to BMS, suggesting intrinsic resistance to RA withdrawal. This difference is consistent with the idea that in AJ lungs, endogenous RA normally constrains TGFβ activity at the onset of airway SM differentiation, and its removal derepresses TGFβ targets that drive an abnormal SM program and airway remodeling.

Quantitative Trait Locus (QTL) mapping in mice has previously identified loci for AHR that colocalize with genes involved in extracellular matrix organization, growth factor signaling, and SM differentiation (4, 5). Many of these loci are syntenic with human GWAS signals associated with asthma and lung function, suggesting a good agreement of our findings with human genetic data, and that our observations may be mechanistically relevant to human disease. In humans, asthma is a polygenic trait with substantial heterogeneity in morbidity, therapeutic response, and airway remodeling phenotypes. Common variants in *TGFB1*, *PDGFRA,* and ECM-related genes have been linked to asthma severity, airway wall remodeling, and reduced lung function (59, 67–71). Notably, the *TGFB1* promoter polymorphism C-509T (rs1800469) in humans has been linked to increased *TGFB1* transcription and heightened AHR and asthma severity (72). Our findings link these human observations to a potential mechanistic framework in which prenatal micronutrient status, through modulation of RA–TGFβ crosstalk, can tip a genetically poised mesenchymal progenitor population toward a remodeling-prone state. The TGFβ–PDGFRA–TNC axis we identified as a core component of this dysregulated program has strong precedents in developmental and pathological remodeling contexts. PDGFRA is a canonical marker of embryonic lung fibroblasts and airway SM progenitors; lineage tracing demonstrates that *Pdgfra*⁺ cells populate peribronchiolar mesenchyme and can differentiate into contractile airway SM cells during development (40, 56, 57). In disease settings, PDGFRA⁺ fibroblasts are increased in the lungs of asthmatics and contribute to airway wall thickening (58). *Tnc* is a well-established TGFβ–SMAD target gene (62) and can be induced synergistically by PDGF and TGFβ signaling (61). Functionally, TNC promotes cell proliferation, migration, and integrin/PDGFR complex signaling in SM cells, and its expression is elevated in the airways and lavage fluid of patients with severe or refractory asthma (73). Variants near or within *PDGFRA*, such as rs1800810, have been linked to severe nonallergic asthma and enhanced airway remodeling (74). GWAS has also implicated *TNC* in asthma risk, with certain alleles affecting ECM stiffness and influencing bronchial mechanics (75). These convergent lines of evidence point to a shared pathway in which genetic variation affecting the regulation or responsiveness of TGFβ–PDGFRA–TNC signaling modulates susceptibility to structural airway disease. Our transcriptomic analyses also identified *Igf1* as a marker of the cluster 2 mesenchymal cells (which also featured *Tgfbi*). IGF1 is a known mitogen for airway SM cells, promoting proliferation and hypertrophy, being also implicated in airway remodeling in asthma (41). Although we found increased *Igf1* expression in mesenchymal populations of AJ-BMS lungs, this gene was not specifically upregulated in *Tgfbi*⁺ cells, raising the possibility that it may be regulated by RA directly or by an additional mechanism.

Our findings suggest that in genetically susceptible individuals, transient environmental perturbations - such as maternal vitamin A insufficiency – during a critical developmental window could trigger abnormal activation of pathways to permanently reprogram specific progenitor niches in a manner dependent on host genetics, predisposing to abnormal remodeling in response to later life. In this context, the observations reported here illustrate well the broader concepts of the developmental origins of health and disease (DOHaD). From a translational perspective, the identification of a discrete, RA-sensitive, TGFβ -activated progenitor population provides both a mechanistic link between prenatal nutrition and adult airway disease and a potential target for future studies aiming at early-life intervention. If analogous cells exist in the human fetal lung—and given the high conservation of developmental signaling pathways between mice and humans, this is likely—then individuals harboring risk alleles in *TGFB1*, *PDGFRA*, *TNC*, or related RA pathway genes may benefit from personalized nutritional care during pregnancy to ensure adequate micronutrient availability during the critical developmental windows.

In conclusion, our results support a model in which a genetically determined RA-sensitive mesenchymal progenitor niche acts as a developmental checkpoint for ASM initiation. In susceptible genetic backgrounds, transient loss of RA signaling unleashes a hyperactive TGFβ–PDGFRA–TNC program that persists beyond development, structurally remodeling the airway and manifesting functionally as AHR in adult life. This model integrates genetic background, environmental exposure, and developmental signaling into a unified framework to tackle AHR susceptibility in the context of asthma and other chronic pulmonary conditions. Future work integrating single-cell genomics of human fetal lung, functional assays of risk allele impact, and longitudinal clinical cohorts with prenatal nutritional data could validate and extend these findings, ultimately enabling precision prevention strategies for airway disease rooted in developmental biology.

## METHODS

### Mice

A/J and C57BL/6J mice were purchased from The Jackson Laboratory (stock no.000646 and stock no.000664). Animals were housed in a pathogen-free environment. For embryonic developmental staging, the morning a vaginal plug was detected was considered E0.5. The genotypes of embryonic mouse sexes were determined by PCR using genomic DNA isolated from embryonic tails and amnions. Animal sex/genders are indicated in each experiment. We included both male and female sexes in this study. Sex was not considered as a biological variable in the study.

### Maternal administration of BMS493

Pregnant mice were orally administered the inverse pan retinoic acid receptor (RAR) agonist, BMS493 (BMS) (3.75, 7.5, or 15 µg/g of BW/day) (Cat#B6688, Sigma-Aldrich), or corn oil from E9.5 to E14.5, a developmental window that encompasses the initial stages of lung morphogenesis and overall initiation of differentiation. With the exception of the dose-response study (Supplemental Figure 1A-B), all other experiments were performed using the BMS concentration of 3.75 µg/g of BW/day, which did not disrupt organogenesis and allowed postnatal development to proceed to adulthood. Dams were either sacrificed for embryonic lung analyses at E14.5 or allowed to continue on a standard nutritionally complete chow diet (15 IU of vitamin A/g of diet; 5053, irradiated PicoLab^R^ Rodent Diet 20) to deliver pups. These pups were allowed to grow to adulthood under the same chow diet for subsequent adult lung analyses and airway resistance measurements.

### Methacholine challenge and specific airway resistance measurements

Adult mice breathing spontaneously were placed in chambers to assess specific airway resistance using whole-body plethysmography (FinePointe for Non-Invasive Airway Mechanics, Data Science International), following the manufacturer’s instructions. Specific airway resistance was assessed in 16-week-old male mice. Baseline specific airway resistance was measured following exposure to nebulized saline (control 0 mg/ml of methacholine), after which increasing concentrations of nebulized methacholine (6.25, 12.5, 25, and 50 mg/ml) were delivered (Cat#1396364, USP).

### Lung explant culture

E12.5 mouse lungs were cultured for 48 hours at 37℃ in 5% CO_2_ on 6 well Transwell-COL dishes containing 1.5ml BGJb medium (Cat#12591038, Thermo Fisher Scientific), 1% Penicillin-Streptomycin (Cat#15140122, Thermo Fisher Scientific), 0.25mg/ml ascorbic acid 0.025% ascorbic acid (Cat#A92902, Sigma-Aldrich), and 1% FBS (Cat#26140079, Thermo Fisher Scientific), with or without SB431542 (Cat#1614, Bio-Techne) for 48h. Specimens were processed for qPCR or fixed for subsequent immunofluorescence analysis as described below.

### Quantitative reverse transcription PCR

Total RNA was extracted from E14.5 lungs of AJ and B6 mice (Control and BMS493 treatment) using the RNeasy Micro Kit (QIAGEN, Cat#74004). cDNA was generated with the High-Capacity cDNA Reverse Transcription Kit (Thermo Fisher Scientific, Cat#4374966). Quantitative real-time PCR (qRT-PCR) was performed in duplicate on a LightCycler 480 II (Roche, IN) using SYBR Green Master Mix (Roche Cat# 04707516001). The thermal cycle was programmed for 5 minutes at 95°C for initial denaturation, followed by 45 cycles of 10 s at 95°C for denaturation, 20 s at 60°C for annealing, and 30 s at 72°C for extension and quantification. The PCR products were assessed by a melting curve analysis (65 to 95°C). Primer sets, designed with Primer3, were normalized to the housekeeping gene *Gapdh or Actb*. The primers used in this study are listed in Supplemental Table 16.

### Immunofluorescence and confocal microscopy analysis

Lungs from 16-week-old male mice and E14.5 male embryos were fixed in 4% paraformaldehyde in PBS at 4°C overnight. After washing in PBS, samples were processed for paraffin-embedding. IF was performed in tissue sections (6-8 μm) that were blocked with 1% bovine serum albumin (Cat#A9418, Sigma-Aldrich) and 0.5% TritonX-100 (Cat#T8787, Sigma-Aldrich) for 1 hour at room temperature. Primary antibodies were incubated in a solution of 1% bovine serum albumin and 0.5% TritonX-100 at 4°C overnight. Sections were then washed with PBS and incubated with Alexa Fluor-conjugated secondary antibodies (1:300) and NucBlue Live Cell Stain ReadyProbes reagent (DAPI) (1:10) (Cat#R37605, Invitrogen) for 1 hour. After washing, samples were mounted with ProLong Gold antifade reagent (Cat#P36930, Invitrogen). Antigen unmasking was done using Citrate-Based solution (Cat#H-3300, Vector Laboratories) heated in a microwave. The following primary antibodies were used: mouse anti-α-Actin (αSMA) (1:300, Cat#sc-32251, Santa Cruz Biotechnology) and rabbit anti-transgelin/SM22 (1:200, Cat#10493-1-AP, proteintech). The following secondary antibodies were used: donkey anti-mouse (conjugated with Alexa Fluor 647) (Cat#A-31571, Invitrogen) and donkey anti-rabbit (conjugated with Alexa Fluor 568) (Cat#A10042, Invitrogen).

### Morphometric analysis

Confocal microscopy was performed using a Zeiss LSM 710 confocal microscope at 20× magnification. IF images of αSMA and SM22 IF from lungs of control group mice and BMS group mice were analyzed using Aivia AI Image Analysis software (Leica Microsystems). Three proximal adult airways and 2-3 distal embryonic airways were selected from each lung; for each adult airway, 3-5 randomly circled areas were analyzed, while for each embryonic airway, 1 circled area was used. The αSMA or SM22-labeled airway smooth muscle cells in these fields were analyzed. The area (in pixels^2^) and/or the numbers of the αSMA or SM22-positive cells were calculated, and the values were normalized to the length of their basement membranes.

### Identifying retinoic acid receptor binding elements containing open-chromatin regions during lung development

We downloaded E14.5, E15.5, and E16.5 mouse lung ATAC-seq datasets from the ENCODE database. (Mouse ENCODE epigenomic data: PRJNA63471 (76), Gene Expression Omnibus (GEO), National Center for Biotechnology Information (NCBI)). We performed the analysis in accordance with the workflow illustrated in Supplemental Figure 2A to identify candidate RA-regulated genes associated with vitamin A status-dependent lung phenotypes.

To obtain the reproducible open chromatin regions (OCRs), we calculated the irreproducible discovery rate (IDR) of the peaks between the experimental replicates (two per time point) (77), and the peaks with IDR<0.05 were used in this study (IDR-OCRs). The annotation of the IDR-OCRs from their corresponding transcription start site (TSS) was performed and visualized using the ChIPpeakAnno package (78). We defined promoter regions as within <2,000 bp upstream and <1,000 bp downstream from the TSS. The mm10 KnownGene information from the UCSC Genome Browser was used to identify the location of the TSS (79). The parameters for finding overlapping IDR-OCRs between time points using the *findOverlappingPeaks* function of the ChIPpeakAnno package were set to the default settings, where a maximum gap between peaks is -1, and a minimum overlap is 0. The identified IDR-OCRs were further screened to see if they contained RARE (RARa (NR)/K562-RARa-ChIP-Seq (Encode)/Homer (motif 304) motif) using HOMER motif analysis (80).

### Measuring retinoid concentrations of tissues

We purchased seven weeks of A/J and B6 female mice from the Jackson laboratory. After one week of acclimation, we euthanized the animals and collected the liver and lungs for retinol and retinyl ester measurements. All animals were fed the LabDiet JL Rat and Mouse/Irr 6F (5LG4) during the acclimation period. Retinol and retinyl esters were measured by reverse-phase high-performance liquid chromatography (HPLC), as previously reported (81, 82). Retinol and retinyl esters (retinyl oleate, linoleate, palmitate, and stearate) were identified by comparing the retention times and spectral data of the experimental compounds with those of authentic standards. Retinyl acetate (Sigma, MO, USA) served as an internal standard.

### Bulk RNA-seq library construction and analysis

Total RNA was extracted from E14.5 lungs from AJ and B6-exposed embryos from mothers subjected to control or BMS diet (n=4 per group) using RNeasy Micro Kit (QIAGEN, Cat#74004). RNA integrity was assessed by Agilent TapeStation 4150 (Agilent Technologies, CA) and library preparation and sequencing were performed by Novogene Co., Ltd., USA. Following the quality assessment of the sequencing files through FastQC (83) and the trimming of low-quality reads and adapter sequences via Cutadapt (84), the sequences were aligned to the mouse GRCm39 reference genome with GENCODE vM36 gene annotation using the STAR aligner (85). The library quality was evaluated using RSeQC (86). Transcript counts were subsequently analyzed with DESeq2 (87). We defined significantly differentially expressed genes (DEGs) as expression levels at least twice as high or low, accompanied by a false discovery rate-adjusted p-value of less than 0.05. Detailed statistics on sequencing and alignment are provided in Supplementary Table 15.

### Transcription Factor Activity Analysis

Transcription factor (TF) activity was assessed using the uncertainty-aware linear model (ULM) approach implemented in the decoupleR package (88). We used the DESeq2 processed bulk RNA-seq datasetas described above. We utilized the Wald test statistic values for our enrichment analysis. The CollecTRI mouse regulatory network (89) was employed as the reference for TF target interactions. Prior to activity inference, we evaluated the overlaps between network target genes and our expression dataset, confirming sufficient coverage for reliable analysis, and then, performed differential expression analysis between experimental conditions (BMS treatment versus control). We selected the entry with the lowest p-value for which multiple Ensembl IDs map to the same gene symbol. A significance threshold of adjusted p-value < 0.05 was applied to extract the significant TF activities.

### Single-nucleus Multiome (snMultiome) sequencing library preparation

We assessed gene expression and chromatin accessibility at single-cell resolution using the 10XGenomic single-cell Multiome platform on E14.5 male mouse lungs of control and BMS493 treatment groups in AJ and B6 strains. To collect enough single nuclei, embryonic lungs from four (B6) or six (AJ) mice were pooled and mechanically dissociated using the gentleMACS Octo Dissociator (Miltenyi Biotech, MD) with the “4C _nuclei_1” program. After dissociation, the sample was incubated on ice for 5 minutes, filtered through a 30 µm filter to remove larger particles, and centrifugated at 4°C 300xg for 5 minutes. The pellet was suspended in 500 µl of cold resuspension buffer (1X PBS, 0.1% BSA, 0.2 U/µl RNase inhibitor), centrifugated at 4°C 300xg for 5 minutes and the pellet was resuspended in 300 µl of cold resuspension buffer. To further purify and concentrate the nuclei, 300 µl of nuclei suspension was gently mixed with an equal volume of 50% OptiPrep (MilliporeSigma, MA) in a 2 ml tube and applied gently on top of a 29% OptiPrep (600 µl) cushion in another 2 ml tube, followed by centrifugation at 4°C 10000xg for 30minutes. The OptiPrep supernatant was poured out in a single motion, and the pellet was resuspended in 0.1X lysis buffer (0.1X nuclei extraction buffer with 0.2 U/µl RNase inhibitor) for permeabilization according to the 10X Genomics nuclei permeabilization step (CG000375). Nuclei were then collected and resuspended in Diluted Nuclei Buffer (1X Nuclei Resuspension Buffer, 1mM DTT, and 1 U/µl RNase inhibitor). Single nuclei were counted with AO/PI staining using a DeNovix CellDrop FL (D.A.I. Scientific Equipment, IL). The integrity of isolated nuclei was inspected under a microscope at 100x magnification. The Multiome libraries were prepared according to the 10X Genomics Chromium Next GEM Single Cell Multiome ATAC +Gene Expression user guide (CG000338). The library quality was evaluated using the Agilent 4200 TapeStation (Agilent Technologies, CA), and the library concentration was determined by the Qubit Fluorometer (Invitrogen, CA). The single-nuclei ATAC-seq (snATAC-seq) and single-nuclei RNA-seq (snRNA-seq) libraries were sequenced on the Illumina NextSeq 2000 using the 50-8-24-49 and 28-10-10-90 formats, respectively, aiming for 25000 snATAC-seq and 20000 snRNA-seq reads per cell to ensure data coverage.

### snMultiome data processing, analyses, and data visualization

#### Preprocessing

Raw FASTQ files were processed using the Cell Ranger ARC pipeline (version 2.0.2, 10x Genomics) to align reads to the mm10 reference genome (2020-A release). This process yielded unique molecular identifier (UMI) counts for gene expression, and ATAC-seq fragment counts for chromatin accessibility. Subsequent analyses were conducted using a standardized snMultiome workflow implemented in R (version 4.4.1), leveraging the Seurat (version 5.0.1) (90–93) and Signac (version 1.14.0) packages (94). Seurat objects were created using the matrix.h5 and fragment.tsv.gz files annotated with EnsDb.Mmusculus.v79. ATAC-seq peaks were identified using MACS2 (ver. 2.2.7.1) (95) and quantified counts in each peak using the FeatureMatrix function in Signac, which was used for downstream analyses of chromatin accessibility. Chromosome nomenclature was standardized to the UCSC style to ensure compatibility with the Cell Ranger ARC output. Quality control measures were applied to exclude low-quality cells from the analysis. Cells were filtered based on the following criteria: low total RNA counts (nfeature_RNA >300, except AJ control >750), low total ATAC-seq counts (nfeature_ATAC>300), high mitochondrial gene expression percentage (<0.2), nucleosome signal > 2, and transcription start site (TSS) enrichment score < 1. These stringent filters ensured the retention of high-quality cells for downstream analyses. The resulting dataset provides a comprehensive view of the transcriptome and chromatin accessibility landscape at single-cell resolution, enabling in-depth exploration of cellular heterogeneity and regulatory mechanisms in the ABC and MNC strains under control and treated conditions.

#### Gene expression data analysis

Gene expression data were analyzed using Seurat (v5.0.1) (93). Quality control was performed by filtering cells based on total RNA counts, percentage of mitochondrial genes, and number of features detected. The remaining cells were normalized using the *NormalizeData* function with default parameters. Highly variable genes (n=3,000) were identified using the *FindVariableFeatures* function, and data were scaled using *ScaleData*. To integrate data across all experimental conditions, we employed Seurat’s integration workflow. Anchor cells were identified between datasets using *FindIntegrationAnchors* (dimensions = 1:30), followed by data integration using *IntegrateData* (dimensions = 1:30). Principal component analysis (PCA) was performed on the integrated data using *RunPCA*. Uniform Manifold Approximation and Projection (UMAP) dimensionality reduction was then applied using *RunUMAP*.

#### ATAC-seq data analysis

ATAC-seq data were processed using Signac (v1.14) (94). Feature selection was performed using *FindTopFeatures*, retaining peaks present in at least 50 cells. Data were normalized using the Term Frequency-Inverse Document frequency (TF-IDF) implemented in *RunTFIDF*. Dimensional reduction was performed using singular value decomposition (SVD) via *RunSVD*. ATAC-seq data were integrated across experimental conditions using the latent semantic indexing (LSI) algorithm in Seurat (96). LSI components were created for each sample independently and then integrated using *FindIntegrationAnchors* and *IntegrateEmbeddings*.

#### Multimodal integration and clustering

To leverage both gene expression and chromatin accessibility data, we employed Seurat’s weighted nearest neighbor (WNN) approach. The *FindMultiModalNeighbors* function was used to compute a neighbor graph, assigning equal weights to RNA-seq and ATAC-seq modalities (weight = 0.5 each). Using the WNN graph, we performed Uniform Manifold Approximation and Projection (UMAP) for visualization and Leiden clustering for cell type identification. The *FindClusters* function was used with a resolution parameter of 0.5, and we specifically sub-clustered a few of the clusters for better annotation and identification using *FindSubCluster*. Clusters were annotated based on the expression of known cell type-specific markers and results from the *FindAllMarkers* function using the Wilcoxon Rank Sum test.

#### Differentially accessible regions (DARs) and differentially expressed genes (DEGs)

Differential gene expression (DEG) and differentially accessible region (DAR) analyses were conducted separately for AJ and B6 strain samples. The *FindMarkers* function in Seurat (93) was used to compare control and treated samples. For DEG analysis, we employed the MAST algorithm (97). For DAR analysis, we used the likelihood ratio (LR) test with no specific thresholds. A significance threshold of adjusted p-value < 0.05 and average log2FC >0.58 and –0.58 was applied to extract the significant differentially expressed genes and peaks for the downstream analysis. We identified the genes closest to the DARs using the *ClosestFeature* function in Signac.

#### Peak-to-gene linkage analysis

We computed the GC content for each peak using the *RegionStats* function in Signac (94), utilizing the BSgenome.Mmusculus.UCSC.mm10 package as the reference genome. Subsequently, we employed the *LinkPeaks* function to infer peak-to-gene linkages, restricting our analysis to the differentially expressed genes identified in our previous analyses.

### Statistical analyses

All statistical analyses were performed using R version 4.1.1 (https://www.r-project.org/). The Student’s *t*-test was used to analyze continuous variables, and Fisher’s exact test was used to analyze categorical variables between control and BMS-treated groups unless otherwise stated. P-values of <0.05 or FDR-adjusted p-values in multiple testing analyses were considered significant.

### Study approval

All experiments conducted on mice were approved and carried out in accordance with the ethical guidelines of the Columbia University Institutional Animal Care and Use Committee (WVC, IACUC #: AC-AABF2567) or the Albert Einstein College of Medicine Institutional Animal Care and Use Committee (MS, IACUC# 00001232) and ARRIVE guidelines.

## Data availability

All data used in the figures are represented in the Supporting Data Values file available online. Raw data of the scRNA-seq analysis performed for this study are available at the Gene Expression Omnibus database under accession number GSE307765. The processed data and code used in this study are available at https://github.com/SuzukiLabTAMU/prenatal_VA_Otoshi.

## AUTHOR CONTRIBUTIONS

Conceptualization and methodology, W.V.C. and M.S.; investigation, T.O., Z.C., Y.S., A.K.S.K., B.J.K., Y.K, P.R., Y.M., S.M.S., and M.S.; formal analysis and software, T.O., Z.C., A.K.S.K., B.J.K., and M.S.; writing—original draft, M.S., W.V.C.; writing—review and editing, all authors; Supervision, L.Q., W.V.C., and M.S.

## ACKNOWLEDGMENTS

The authors thank Dr. Jun Fan and the team members of the Molecular Genomics core at Texas A&M Institute for Genome Science and Society for their help in single nucleotide Multiomics library preparations. Portions of this research were conducted with the advanced computing resources provided by Texas A&M High Performance Research Computing.

## SUPPLEMENTAL FIGURE LEGENDS

**Supplemental Figure 1. Differential effects of prenatal disruption of RA signaling by maternal BMS-containing diet in AJ and B6 mice.**

**(A)** Diagram experimental design: maternal administration of control (corn oil) or BMS-containing diet from gestation day 9.5-14.5 to AJ and B6 adult mice.

**(B)** Effect of maternal administration of a diet containing different BMS concentrations (3.75-15 μg/body weight per day (μg/bw/day) on the gross morphology of E14.5 embryos and lungs in both strains. Body truncation and lung hypoplasia at the highest maternal BMS concentration most prominent in AJ embryos compared to B6. These effects were not seen at 3.75 μg/bw/day in either strain. Both controls (corn oil) and BMS (3.75 μg/bw/day) pups reached adulthood.

**(C)** Expression of RA pathway components (*Rarb*, *Cyp26b1*, *Lrat*, *Stra6*) and *Epcam* in lung homogenates from E14.5 AJ and B6 embryos (qPCR). Graphs: No downregulation in *Epcam* expression between control and BMS-exposed lungs in spite of significant *Rarb*, *Cyp26b1*, *Lrat, Stra6* downregulation by BMS in both strains. *Cyp26b1*, *Lrat and Stra6* expression significantly higher in control AJ compared to control B6, while *Rarb* is higher in control B6. \**p*<0.05. \*\**p*<0.01.

**(D)** Bulk RNA-seq analysis (*Lrat, Stra6, Dhrs3)*: whole E14.5 lungs from AJ and B6 embryos exposed to maternal BMS (3.75ug/g/bw) diet: downregulation by BMS in both strains and confirming the increased *Stra6* and *Lrat* expression in control AJ compared to B6 control.

**(E)** Effects of prenatal BMS exposure (3.75ug/g/bw) in adult AJ and B6 offsprings: body weight, lung and liver weight relative to body weight. Graphs are mean ± SE, \**p*<0.05. AJ Control (n=6), AJ BMS (n=6), B6 Control (n=9), B6 BMS (n=6).

**Supplemental Figure 2. Differential expression of RA-responsive genes in embryonic lungs from AJ and B6 mice underlies the basis for the functional differences between these strains.**

**(A)** Schematic overview: workflow for identification of RA–responsive open chromatin regions (RARE-OCRs) enriched in the embryonic lung during early differentiation (mouse ENCODE ATAC-seq datasets epigenomic: PRJNA63471) (see Methods). Venn diagram (left): overlap of IDR-OCRs in embryonic mouse lungs. A total of 34,039, 26,128, and 45,988 IDR-OCRs were identified at E14.5, E15.5, and E16.5, respectively, with 22,812 regions shared across all three timepoints. While most IDR-OCRs were detected at all three time points, we defined RARE-OCRs as those that were detected as IDR-OCRs at two or more time points. Pie chart (right): Among 28,355 RARE-OCRs, 44.2% were identified within the promoter regions of genes.

**(B)** Bulk RNAseq analysis of E14.5 AJ vs B6 (controls only) identifies 1258 DEGs and strain-specific functional pathways potentially regulated through RARE-containing genes. Volcano plot highlighting *Lrat* upregulated in AJ. Gene Ontology molecular function enrichment analysis of differentially expressed genes containing RARE motifs. Bar plots: significantly enriched GO terms (padj < 0.05) for top RARE-OCR genes in each strain (AJ-high genes and B6-high genes), ranked by significance.

**(C)** Bulk RNAseq of BMS vs control E14.5 AJ and B6 lungs. Left: Venn diagram indicating the number of DEGs downregulated or upregulated by BMS from each strain. DEGs downregulated in both strains include markers of RA activity (*), genes harboring RARE-OCR (bold) and DEGs in controls (red: AJ; blue: B6). Right: Hox family member genes downregulated in BMS-exposed lungs from both AJ and B6 embryos. Graphs: n=4 per group.

**(D)** Gene Set Enrichment Analysis (GSEA) of DEGs in BMS vs control E14.5 lungs from B6 mice. Dot plots: significantly enriched Gene Ontology terms for biological processes (BP) and molecular functions (MF) displayed by direction of enrichment (activated vs. suppressed pathways). GO terms enriched in AJ BMS include suppression of RA-related molecular function (RA binding) and activation of biological processes related to muscle contraction and microfilament activity (Figure 2E). No molecular function GO terms were detected in B6, suggesting a less robust RA-related functional activity.

**Supplemental Figure 3. Multiome analysis reveals significantly higher number of DEGs and open chromatin regions in AJ than B6 lungs in response to intrauterine RA signaling disruption.**

**(A)** Split UMAP projections displaying the distribution of the four major cell populations across conditions (control and BMS in AJ and B6 lungs).

**(B)** Volcano plots showing differentially expressed genes (DEGs) and differentially accessible chromatin regions (DARs) in epithelial, mesenchymal, and endothelial cells in response to BMS in AJ and B6 lungs. Significantly higher number of DEGs and DARs in AJ compared to B6. Mesenchymal compartment showing the greatest number of DEGs/DARs (boxes).

**Supplemental Figure 4. Distinct subpopulations of ECM-producing, vascular and airway SM precursors in the mesenchymal compartment of developing lung.**

**(A)** U-map (left) overview cluster analysis of mesenchymal compartment from all groups. Cluster 0 (undifferentiated proliferative) signature characterized by cell cycle-related genes (*Top2a, Kif15, and Mki67*), visualized by dot plot (top right), violin plots, and feature plots. In situ hybridization (ISH) showing diffuse transcript distribution throughout the E14.5 lung mesenchyme.

**(B-C)** Dot plots: enrichment of mitochondrial genes in cluster 4 and extracellular matrix-associated genes in cluster 6.

**(D-E)** Dot plot of top Cluster 5 genes featuring *Pdgfrb, Heyl, Notch3, Cspg4* identified by their association with the vascular smooth muscle (SM) program. Violin plots, feature plots and ISH images of representative cluster 5 genes. Venn diagram showing distinct and shared gene expression of clusters 5 (vascular) and 7 (airway) SM populations. Images depict striking difference in spatial distribution of *Myom1* (cluster 7) and *Pdgfrb, Heyl, Notch3* (cluster 5) reflecting the distinct airway and vascular SM signatures in the E14.5 lung mesenchyme.

**Supplemental Figure 5. Multiome analysis identifies a distal mesenchymal population enriched in regulators of cell migration and tissue patterning genes.**

**(A)** Top differentially enriched genes of Cluster3 (left). Table: most significant terms ranked by adjusted p value (Enrichr, Descartes Cell Types and Tissue 2021 library). Terms in red are lung-related. Violin and feature plots of representative cluster 3 genes (*Sema3d, Epha3 and Ptn)*. ISH showing most distal expression in the subpleural mesenchyme.

**(B and C)** Table depicting additional components of the Cluster 3 signature (left) featuring key regulators of lung pattern formation and mesenchymal differentiation (right: GO enrichment bar plot and terms listing these components). Genes shown in red in the tables are representative marker genes for Cluster 3. Gene ontology (GO) enrichment analysis reveals strong association with cell migration, morphogenetic movements and chemoattraction reported mostly in neuronal studies. *Wnt2* and *Fgf10* identified in Cluster 3 by dot plot, violin plot and feature plot. ISH of *Wnt2, Fgf10, Rspo2, Cttna2* showing distal mesenchymal localization, including subpleural region.

**(A) (D)** Proposed developmental stages of airway SM differentiation based on our data and a model previously reported (Kumar et al. 2014). A three-stage progression from cluster 3 (undifferentiated tip bud mesenchymal progenitors) and cluster 0 (proliferative progenitors), through cluster 2 (recruitment zone), culminating in mature ASM in cluster 7 (maturation zone).

**Supplemental Figure 6. Differentially expressed genes between AJ and B6 control E14.5 lungs reflect strain differences in genetic background.**

**(A)** Volcano plots: greater number of DEGs in cluster 0 (478), cluster 2 (273), and cluster 3 (161), with fewer DEGs detected in the remaining clusters. The total number of cells per cluster is indicated. Significantly upregulated or downregulated genes (|log2FC| > 0.58, padj < 0.05) are represented in red.

**(B)** *Rarb* expression across mesenchymal clusters in AJ and B6 controls (dot plots and violin plots). No statistically significant difference in *Rarb* between AJ and B6 in any cluster.

**(C-F)** Identification of differentially upregulated and downregulated genes between control AJ and B6 lungs in clusters 0, 2, and 3. Heatmaps and table of the gene expression signature of individual clusters (**C,D)** and whole lung mesenchyme (clusters 0-9). The total cells per strain per cluster are shown in the heatmaps. **(E)** reveals common genes emerging as top DEGs across all mesenchymal clusters (Ex. *Mgp, Negr1, Hmga2, Col1a1, Opcml*) suggesting a shared transcriptional signature that distinguishes AJ from B6 mesenchymal cells. Several of these genes were also identified as significantly enriched in control AJ or B6 by bulk RNA-seq (whole lung), highlighting concordant strain-specific gene expression differences **(F).**

**Supplemental Figure 7. Transcriptomic responses of *Tgfbi*+ cells to BMS in AJ and B6 lungs.**

**(A)** Markedly different number of DEGs in *Tgfbi*+ cells between control and BMS-exposed AJ (top) and B6 (bottom) lungs.

**(B)** Identification of cluster 2-enriched genes among those downregulated by BMS from whole-mesenchymal analysis of AJ and B6 lungs. Data represent the log₂(fold change) in expression between BMS and control across mesenchymal clusters (x-axis) against the log₂(fold change) of cluster 2 genes (y-axis).

**(C)** Identification of cluster 2-enriched genes among the DEGs upregulated by BMS in *Tgfbi*+ cells of B6 lungs (adjusted p-value <0.05 and log2 Fold Change > 0). Plot represents the log₂(fold change) in expression between BMS and control conditions in *Tgfb*i+ cells (x-axis) against the log₂(fold change) of cluster 2 genes (y-axis). None of the DEGs found carried Smad2, Smad3, or Smad4 binding sites.

