## Supplemental Figures for "Genetic background and transient prenatal disruption of vitamin A signaling determine susceptibility to airway hyperresponsiveness in mice"

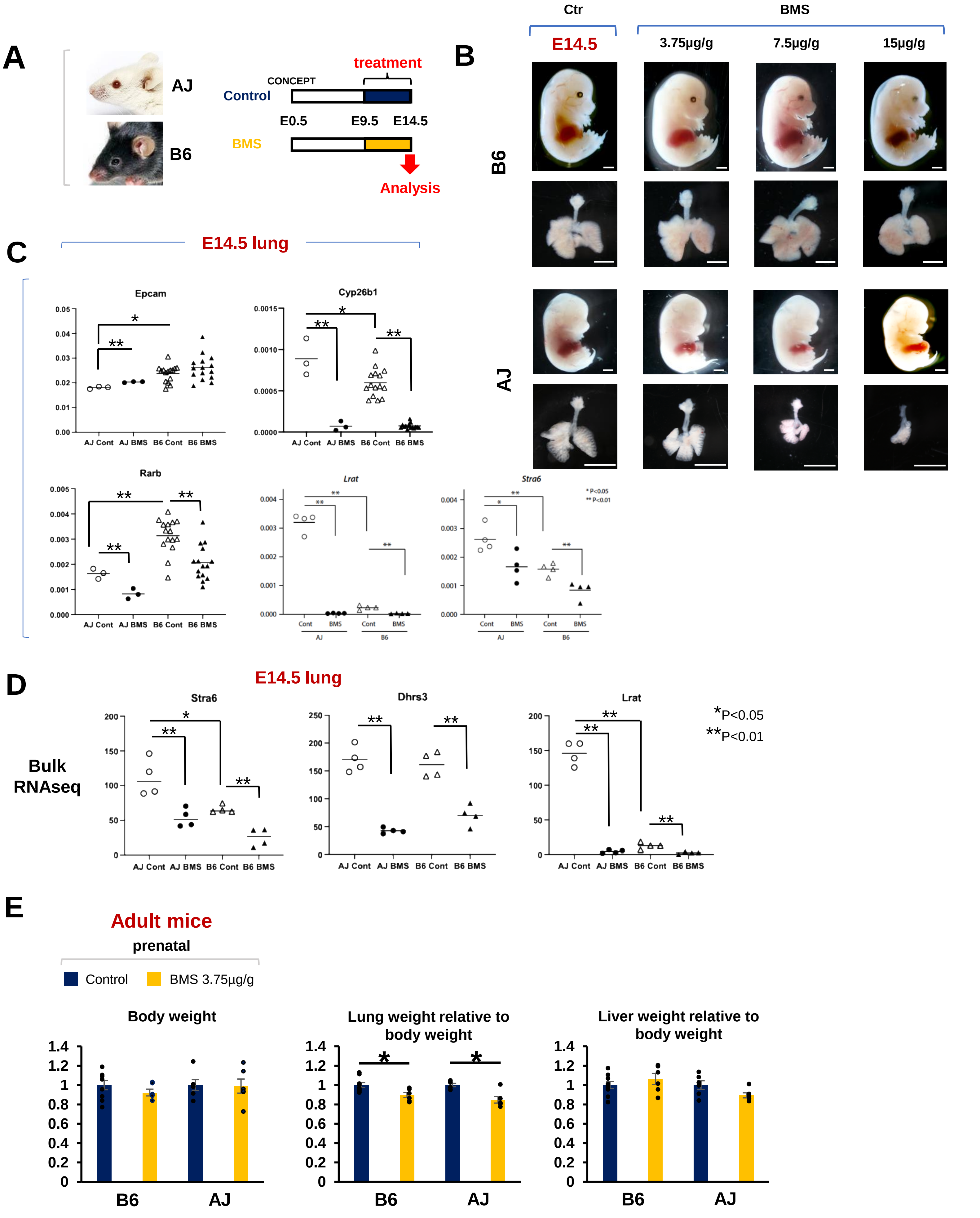


Suppl. Figure 1
Otoshi et al.

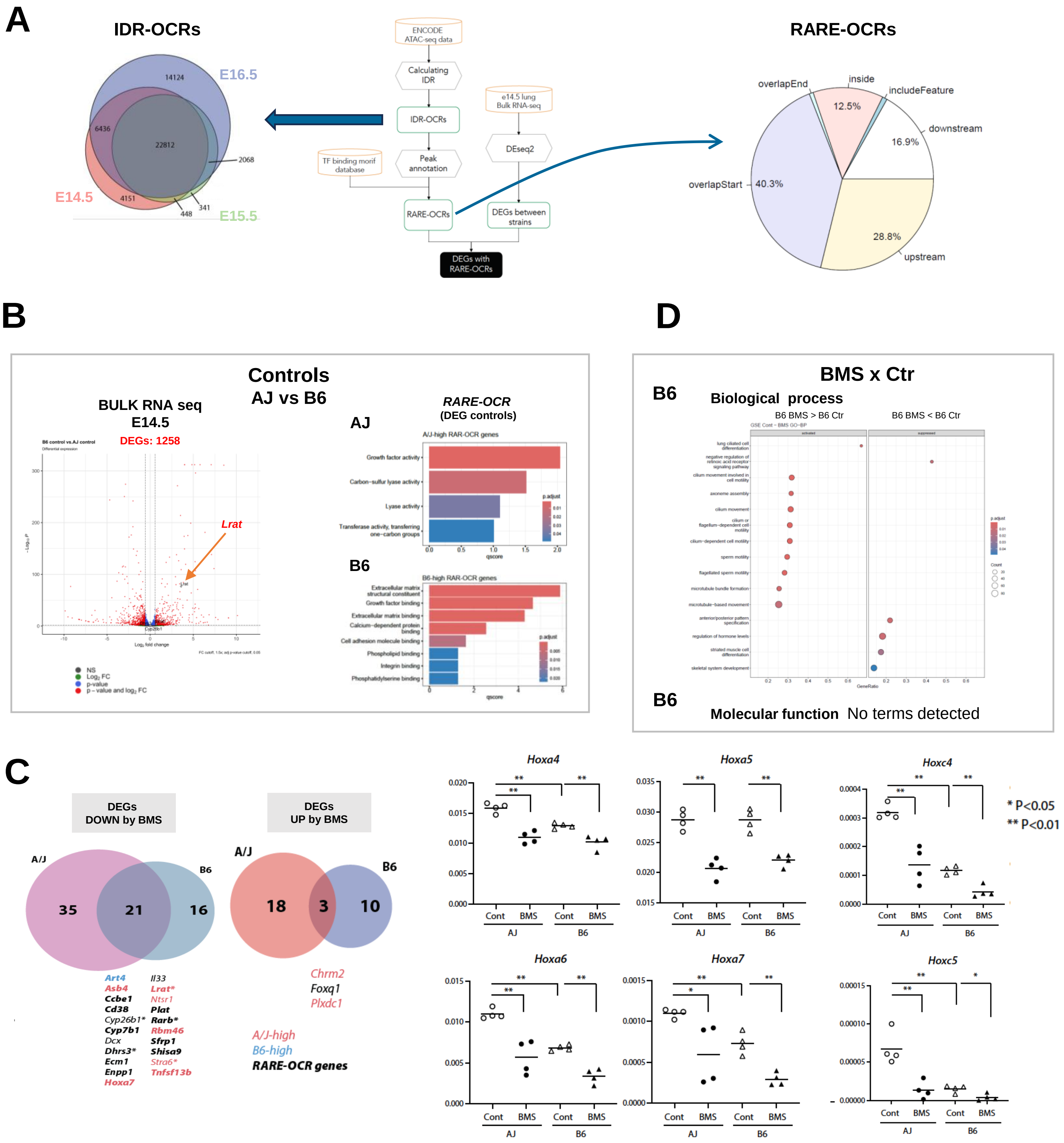


Suppl. Figure 2
Otoshi et al.

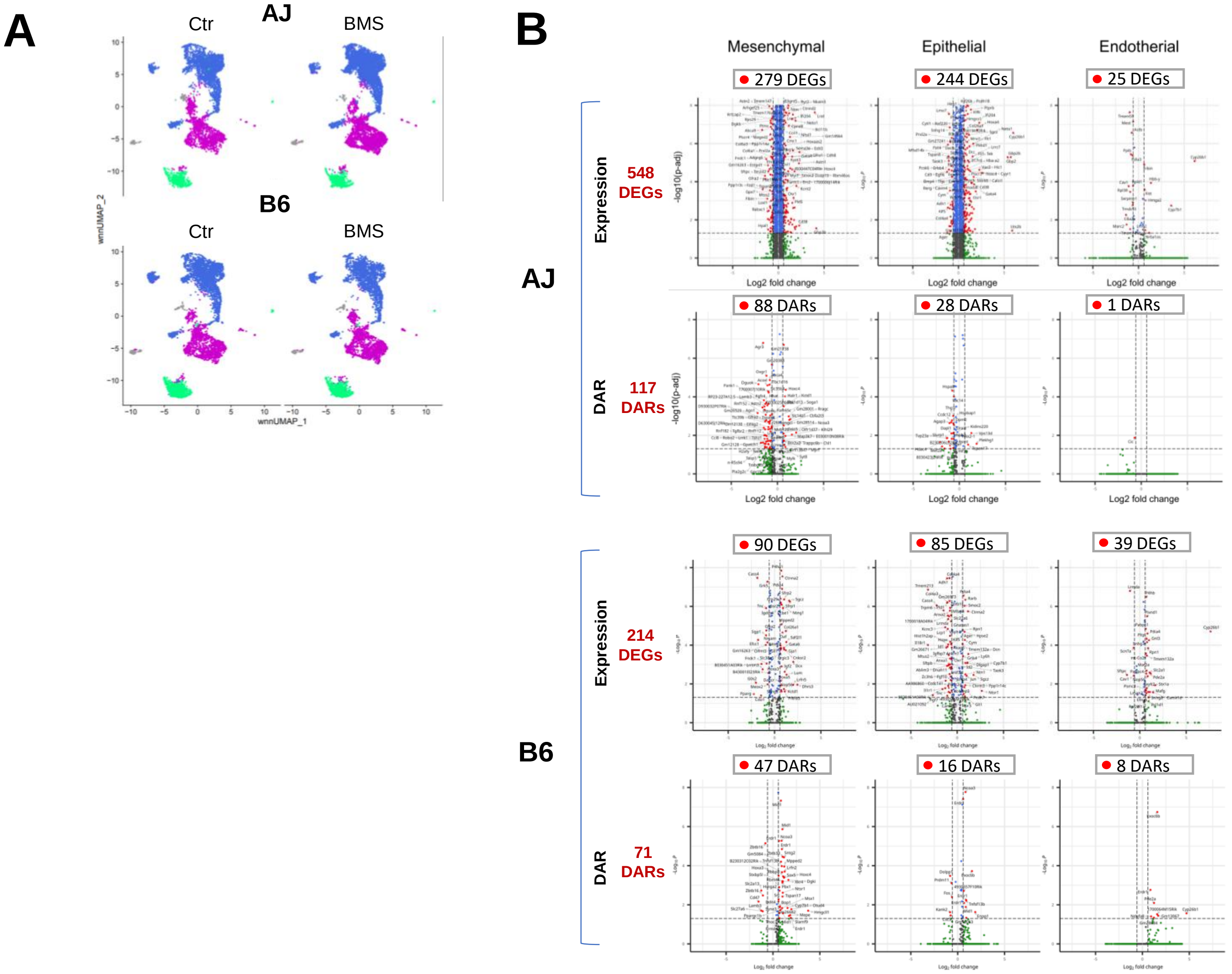


**Suppl. Figure 3**
**Otoshi et al.**

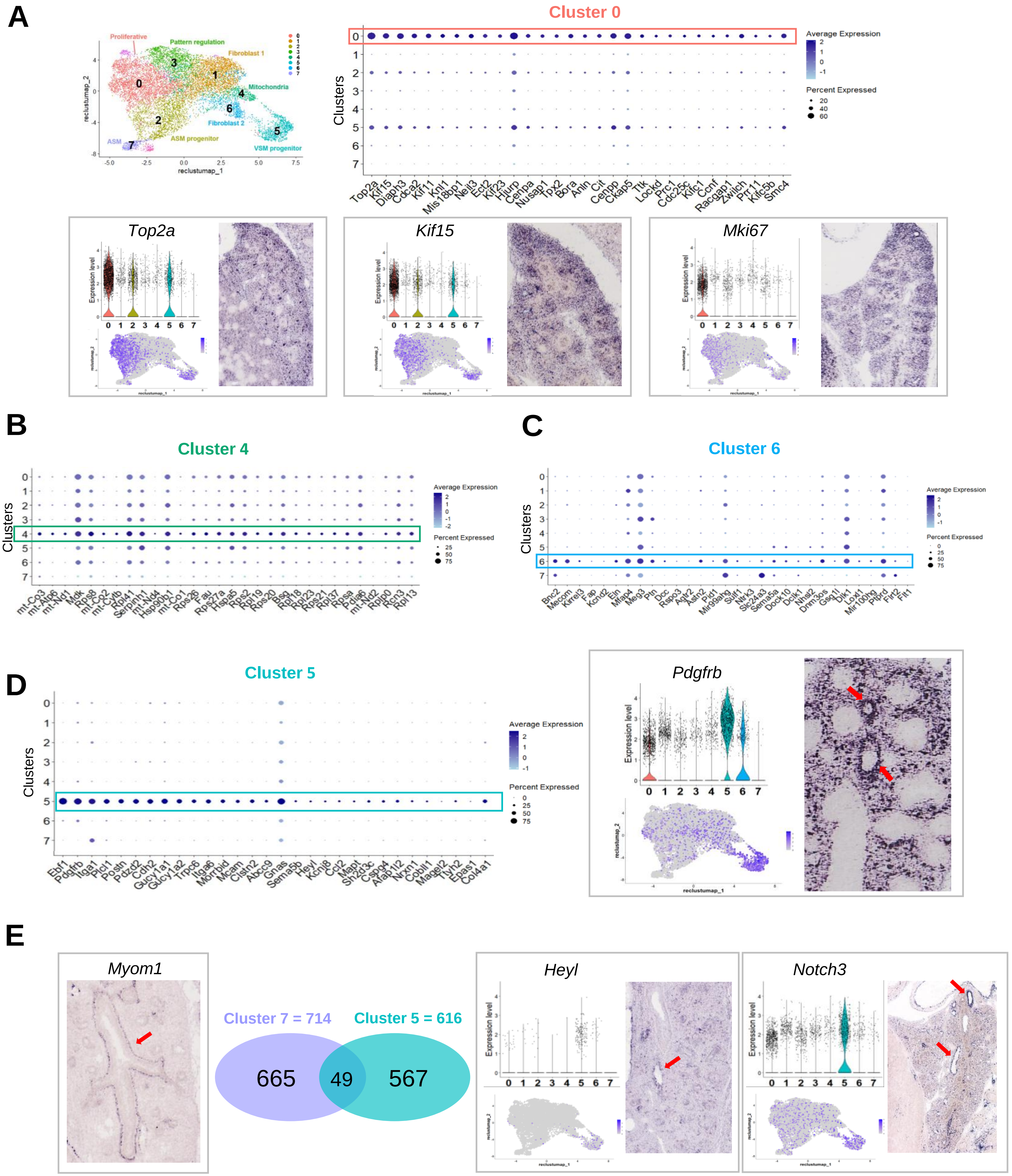


Suppl. Figure 4
Otoshi et al.

**A**

### Cluster 3

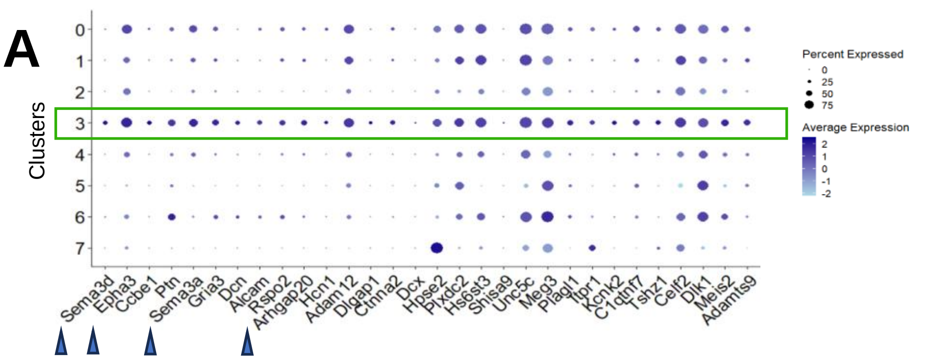


**Descartes Cell Types and Tissue**

| Name | Adjusted p value |
|---|---|
| Visceral neurons in Lung | 0.000006813 |
| ENS neurons in Stomach | 0.0006165 |
| Stromal cells in Thymus | 0.0008421 |
| Stromal cells in Kidney | 0.001193 |
| Stromal cells in Heart | 0.009271 |
| Stromal cells in Adrenal | 0.009271 |
| Stellate cells in Liver | 0.009271 |
| Stromal cells in Lung | 0.01401 |
| Vascular endothelial cells in Thymus | 0.02486 |
| Stromal cells in Muscle | 0.03563 |

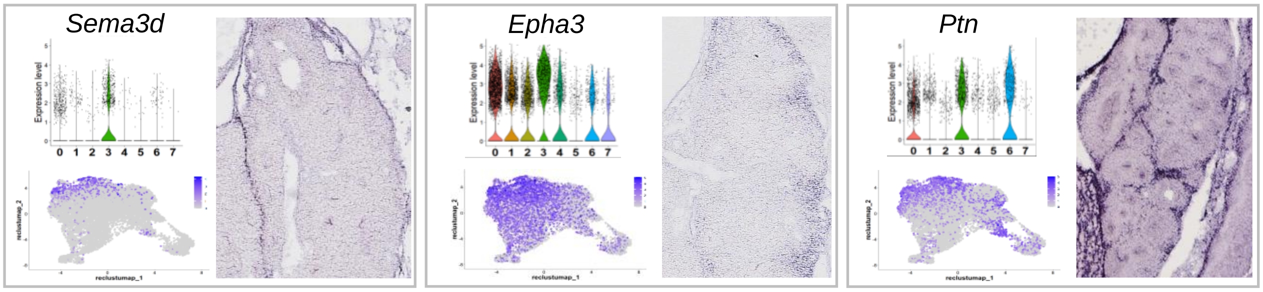


**B**

| Gene | AvgLog2FC | Adj p value ↑ |
|---|---|---|
| Rspo2 | 1.52 | $6.6\times10^{-75}$ |
| Arhgap20 | 1.61 | $6.9\times10^{-75}$ |
| Hcn1 | 2.17 | $1.1\times10^{-74}$ |
| Adam12 | 0.94 | $1.7\times10^{-74}$ |
| Dlgap1 | 2.54 | $1.2\times10^{-67}$ |
| Ctnna2 | 1.78 | $4.4\times10^{-65}$ |
| Dcx | 3.28 | $1.2\times10^{-64}$ |
| Hpse2 | 0.97 | $9.1\times10^{-64}$ |
| Plxdc2 | 0.91 | $8.6\times10^{-59}$ |
| Hs6st3 | 0.77 | $4.5\times10^{-57}$ |
| Shisa9 | 2.61 | $2.7\times10^{-56}$ |
| Unc5c | 0.53 | $4.2\times10^{-56}$ |
| Meg3 | 0.70 | $1.6\times10^{-51}$ |
| Plagl1 | 1.36 | $3.1\times10^{-49}$ |
| Itpr1 | 1.38 | $5.5\times10^{-49}$ |
| Kcnk2 | 1.66 | $5.4\times10^{-47}$ |
| C1qtnf7 | 1.05 | $9.2\times10^{-47}$ |
| Tshz1 | 1.39 | $1.3\times10^{-45}$ |
| Celf2 | 0.61 | $2.9\times10^{-45}$ |
| Dlk1 | 0.63 | $8.1\times10^{-44}$ |
| Meis2 | 1.01 | $5.1\times10^{-43}$ |
| Adamts9 | 1.07 | $8.1\times10^{-43}$ |
| Itga8 | 0.98 | $1.8\times10^{-41}$ |
| Slit3 | 0.69 | $1.5\times10^{-40}$ |
| Fgf10 | 1.17 | $2.6\times10^{-39}$ |
| Capn6 | 1.86 | $4.0\times10^{-39}$ |
| Rian | 0.87 | $4.1\times10^{-38}$ |
| Wnt2 | 1.21 | $2.3\times10^{-35}$ |

### GO enrichment Cluster 3

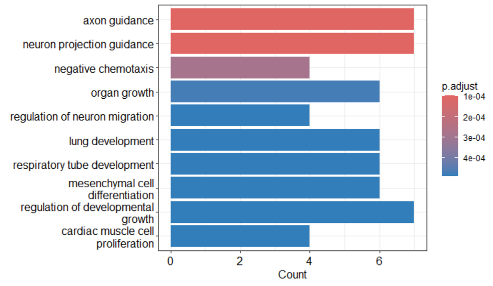


| GO term | Genes |
|---|---|
| axon guidance | *Sema3d*, *Epha3*, *Sema3a*, *Alcam*, *Unc5c*, *Slit3*, *Epha7* |
| lung development | *Ccbe1*, *Rspo2*, *Meg3*, *Fgf10*, *Wnt2*, *Gata6* |
| mesenchymal cell differentiation | *Sema3d*, *Epha3*, *Sema3a*, *Fgf10*, *Rian*, *Wnt2* |

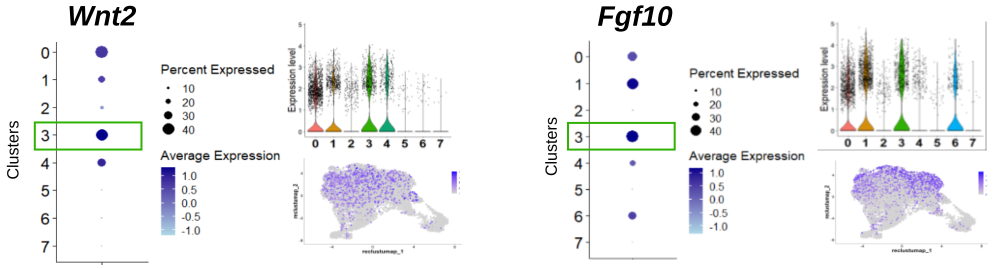


**C**

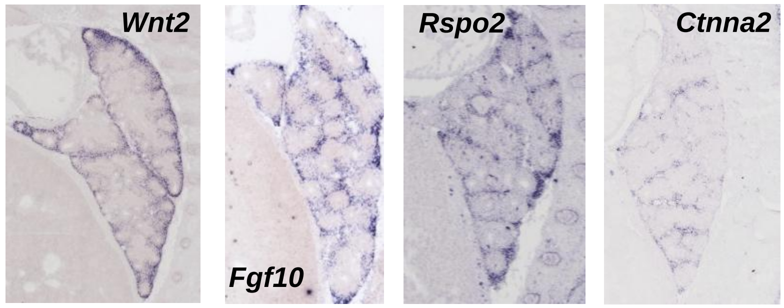


**D**

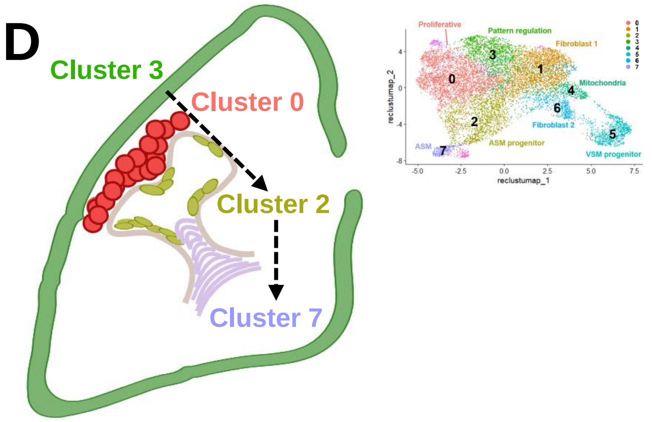


Suppl. Figure 5
Otoshi et al.

**A**

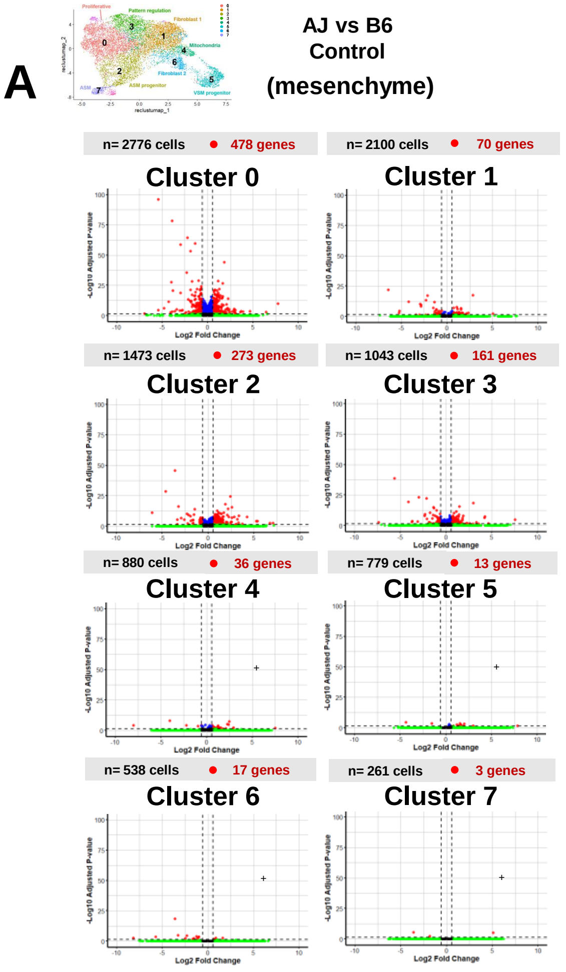


**B**

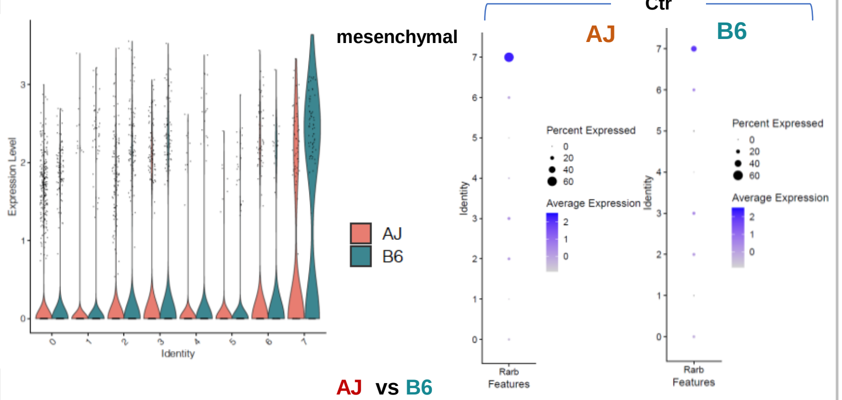


AJ vs B6
Control

**Cluster 0**

| | avg_log2FC | pct.1 | pct.2 | p_val_adj |
|---|---|---|---|---|
| Rarb | 0.37084952 | 0.209 | 0.132 | 0.259067 |

**Cluster 1**

| | avg_log2FC | pct.1 | pct.2 | p_val_adj |
|---|---|---|---|---|
| Rarb | -0.343422 | 0.035 | 0.055 | 1 |

**Cluster 2**

| | avg_log2FC | pct.1 | pct.2 | p_val_adj |
|---|---|---|---|---|
| Rarb | -0.4443896 | 0.217 | 0.176 | 1 |

**Cluster 3**

| | avg_log2FC | pct.1 | pct.2 | p_val_adj |
|---|---|---|---|---|
| Rarb | 0.1572566 | 0.239 | 0.169 | 1 |

**Cluster 4**

| | avg_log2FC | pct.1 | pct.2 | p_val_adj |
|---|---|---|---|---|
| Rarb | -0.39676 | 0.101 | 0.068 | 1 |

**Cluster 5**

| | avg_log2FC | pct.1 | pct.2 | p_val_adj |
|---|---|---|---|---|
| Rarb | 0.6644587 | 0.085 | 0.05 | 1 |

**Cluster 6**

| | avg_log2FC | pct.1 | pct.2 | p_val_adj |
|---|---|---|---|---|
| Rarb | 0.8317866 | 0.25 | 0.149 | 1 |

**Cluster 7**

| | avg_log2FC | pct.1 | pct.2 | p_val_adj |
|---|---|---|---|---|
| Rarb | -1.137191 | 0.349 | 0.648 | 1 |

**C**

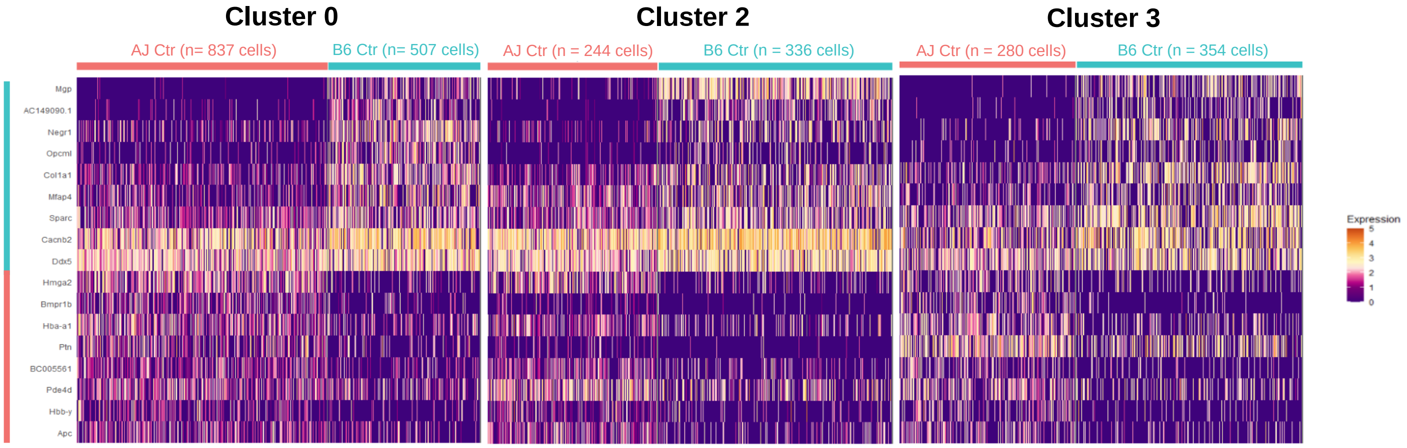


**D**

**Cluster 0**

| Gene | AvgLog 2FC | Adj p value ↑ |
|---|---|---|
| *Mgp* | -5.44 | $7.8\times10^{-97}$ |
| *AC149090.1* | -3.92 | $2.9\times10^{-79}$ |
| *Negr1* | -2.20 | $3.0\times10^{-65}$ |
| *Gm47283* | -1.36 | $1.8\times10^{-60}$ |
| *Opcml* | -2.96 | $1.7\times10^{-59}$ |
| *Col1a1* | -1.90 | $3.7\times10^{-54}$ |
| *Hmga2* | 1.85 | $8.2\times10^{-45}$ |
| *Mid1* | -2.30 | $2.4\times10^{-36}$ |
| *Colec12* | -1.01 | $1.7\times10^{-29}$ |
| *Gm49708* | 1.18 | $2.8\times10^{-29}$ |
| *Mfap4* | -1.70 | $4.1\times10^{-29}$ |
| *Myof* | -3.98 | $2.4\times10^{-28}$ |
| *Bmpr1b* | 1.66 | $2.1\times10^{-27}$ |
| *Nisch* | -0.70 | $7.9\times10^{-25}$ |
| *Ptprd* | -0.69 | $3.6\times10^{-23}$ |
| *Hba-a1* | 1.29 | $8.8\times10^{-23}$ |
| *Slit2* | -1.14 | $1.5\times10^{-22}$ |
| *Hjurp* | -0.76 | $1.9\times10^{-22}$ |
| *Sparc* | -0.82 | $2.9\times10^{-22}$ |
| *Elmo1* | -1.89 | $4.6\times10^{-22}$ |
| *Gpc6* | 0.82 | $5.5\times10^{-22}$ |

**Cluster 2**

| Gene | AvgLog 2FC | Adj p value ↑ |
|---|---|---|
| *Mgp* | -3.59 | $1.9\times10^{-46}$ |
| *AC149090.1* | -4.64 | $2.0\times10^{-29}$ |
| *Mpped2* | 2.41 | $3.1\times10^{-25}$ |
| *Hmga2* | 1.88 | $5.2\times10^{-19}$ |
| *Egfem1* | -3.21 | $8.1\times10^{-17}$ |
| *Tmem158* | 2.60 | $3.2\times10^{-16}$ |
| *Sdk1* | 2.19 | $4.4\times10^{-15}$ |
| *Gm47283* | -1.44 | $8.2\times10^{-15}$ |
| *Mir99ahg* | 1.00 | $1.5\times10^{-13}$ |
| *Mtcl1* | 1.16 | $6.4\times10^{-12}$ |
| *Kif26b* | 1.70 | $6.7\times10^{-12}$ |
| *Myof* | -6.11 | $7.2\times10^{-12}$ |
| *Gm15261* | 1.32 | $3.2\times10^{-11}$ |
| *Kcnn2* | 1.48 | $6.6\times10^{-11}$ |
| *Rims1* | -2.31 | $7.2\times10^{-11}$ |
| *Magi1* | 0.73 | $1.7\times10^{-10}$ |
| *Apc* | 1.02 | $1.8\times10^{-10}$ |
| *Prickle1* | 0.86 | $2.4\times10^{-10}$ |
| *Col1a1* | -1.55 | $2.5\times10^{-10}$ |
| *BC005561* | 1.02 | $4.0\times10^{-10}$ |
| *Negr1* | -2.15 | $5.7\times10^{-10}$ |

**Cluster 3**

| Gene | AvgLog 2FC | Adj p value ↑ |
|---|---|---|
| *Mgp* | -5.63 | $1.9\times10^{-39}$ |
| *Negr1* | -2.98 | $7.6\times10^{-24}$ |
| *Col1a1* | -2.13 | $6.2\times10^{-23}$ |
| *AC149090.1* | -4.12 | $3.4\times10^{-20}$ |
| *Bmpr1b* | 3.00 | $4.1\times10^{-19}$ |
| *Gm49708* | 1.48 | $1.9\times10^{-16}$ |
| *Gm47283* | -1.39 | $1.3\times10^{-15}$ |
| *Col3a1* | -1.34 | $4.3\times10^{-15}$ |
| *Sparc* | -1.05 | $3.4\times10^{-13}$ |
| *Gas5* | 1.11 | $3.3\times10^{-12}$ |
| *Dcn* | -1.76 | $4.0\times10^{-11}$ |
| *Hba-a1* | 0.87 | $8.0\times10^{-11}$ |
| *Nisch* | -0.67 | $3.6\times10^{-10}$ |
| *Opcml* | -2.29 | $5.0\times10^{-10}$ |
| *Igfbp5* | 1.42 | $1.3\times10^{-9}$ |
| *Mfap4* | -1.39 | $3.3\times10^{-9}$ |
| *Zfp292* | 0.39 | $6.3\times10^{-9}$ |
| *Foxp1* | 1.86 | $1.1\times10^{-8}$ |
| *Mid1* | -2.32 | $1.3\times10^{-8}$ |
| *Col1a2* | -1.19 | $1.4\times10^{-8}$ |
| *Tenm4* | 0.72 | $1.4\times10^{-8}$ |

**E**

**Whole mesenchyme**

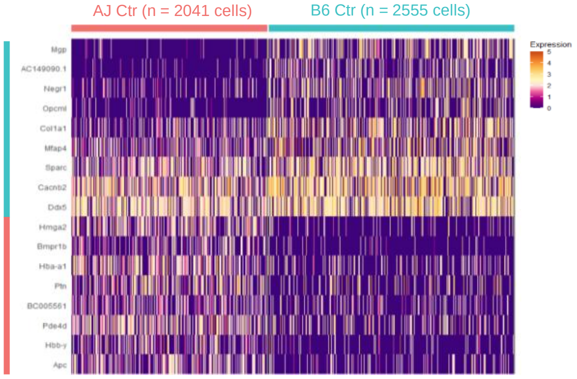


**F**

**AJ Ctr vs B6 Ctr**
**Bulk RNA seq**

| Gene | AvgLog2FC | Adj p value ↑ |
|---|---|---|
| *Mgp* | -2.99 | $4.0\times10^{-251}$ |
| *Opcml* | -2.70 | $2.0\times10^{-96}$ |
| *Negr1* | -2.04 | $3.0\times10^{-72}$ |
| *Ptn* | 0.80 | $3.8\times10^{-32}$ |
| *Mfap4* | -0.73 | $5.9\times10^{-19}$ |
| *Col1a1* | -0.75 | $1.0\times10^{-15}$ |
| *Hmga2* | 1.10 | $5.9\times10^{-13}$ |
| *Hbb-y* | 1.02 | $2.4\times10^{-8}$ |
| *Hba-a1* | 0.64 | $3.2\times10^{-2}$ |

**Suppl. Figure 6**
**Otoshi et al.**

**A**

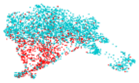

### *Tgfbi+ cells*
(BMS vs Ctrl)
**AJ**

Top 50 (TOTAL = 337)

| Gene | AvgLog2FC | Adj p value ↑ |
|---|---|---|
| *Ddx5* | 0.61 | $3.8\times10^{-17}$ |
| *Hmga2* | -0.98 | $1.6\times10^{-15}$ |
| *Cd63* | 0.91 | $3.2\times10^{-15}$ |
| *Hbb-y* | -2.51 | $1.1\times10^{-13}$ |
| *Epha7* | -0.87 | $1.3\times10^{-12}$ |
| *Unc5c* | -0.59 | $2.5\times10^{-11}$ |
| *Nr6a1os* | -0.64 | $2.7\times10^{-11}$ |
| *Sema3a* | -0.82 | $5.0\times10^{-11}$ |
| *Thsd4* | -0.41 | $1.9\times10^{-10}$ |
| *Hecw2* | -1.24 | $2.3\times10^{-10}$ |
| *Ccbe1* | -2.54 | $2.9\times10^{-10}$ |
| *Pdgfra* | 0.94 | $4.7\times10^{-10}$ |
| *Meg3* | -0.39 | $7.4\times10^{-8}$ |
| *2610307P16Rik* | -0.55 | $9.1\times10^{-8}$ |
| *Spred1* | -0.24 | $1.3\times10^{-7}$ |
| *Prpf40a* | -0.55 | $1.4\times10^{-7}$ |
| *Hsp90b1* | 0.43 | $1.7\times10^{-7}$ |
| *Prex2* | -0.83 | $2.7\times10^{-7}$ |
| *Tnc* | 1.67 | $2.7\times10^{-7}$ |
| *Bmpr1b* | -0.81 | $4.0\times10^{-7}$ |
| *Pdzd2* | -0.67 | $5.0\times10^{-7}$ |
| *Pdia3* | 0.34 | $6.8\times10^{-7}$ |
| *A630089N07Rik* | -0.24 | $8.5\times10^{-7}$ |
| *Ppic* | 0.91 | $9.8\times10^{-7}$ |
| *Ptk2* | -0.42 | $1.0\times10^{-6}$ |
| *Plagl1* | -0.69 | $1.1\times10^{-6}$ |
| *Lrfn5* | -3.29 | $1.9\times10^{-6}$ |
| *Itm2b* | 0.88 | $2.9\times10^{-6}$ |
| *Trim24* | -0.40 | $3.2\times10^{-6}$ |
| *Dnmt3a* | 0.29 | $3.9\times10^{-6}$ |
| *Tshz3* | -0.43 | $5.8\times10^{-6}$ |
| *Ccny* | -0.29 | $5.9\times10^{-6}$ |
| *P3h2* | -1.23 | $7.3\times10^{-6}$ |
| *Mfap4* | 0.65 | $7.5\times10^{-6}$ |
| *Tent2* | -0.44 | $9.8\times10^{-6}$ |
| *Nisch* | 0.40 | $1.0\times10^{-5}$ |
| *Cyp7b1* | -1.09 | $1.2\times10^{-5}$ |
| *Slit3* | -0.51 | $1.3\times10^{-5}$ |
| *Hotairm1* | -1.21 | $1.3\times10^{-5}$ |
| *Hba-a1* | -1.40 | $1.5\times10^{-5}$ |
| *Laptm4a* | 0.39 | $1.7\times10^{-5}$ |
| *Sulf1* | -1.06 | $1.9\times10^{-5}$ |
| *Rbm46* | -0.87 | $2.0\times10^{-5}$ |
| *Dlc1* | -0.29 | $2.1\times10^{-5}$ |
| *Slc25a36* | -0.26 | $3.0\times10^{-5}$ |
| *Tnrc6a* | -0.19 | $3.0\times10^{-5}$ |
| *Frmd4a* | -0.25 | $3.0\times10^{-5}$ |
| *Ptbp2* | -0.46 | $3.1\times10^{-5}$ |
| *Cd81* | 0.39 | $3.5\times10^{-5}$ |
| *Rspo2* | -0.79 | $3.5\times10^{-5}$ |

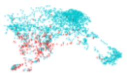

### *Tgfbi+ cells*
(BMS vs Ctrl)
**B6**

TOTAL = 6

| Gene | AvgLog2FC | Adj p value ↑ |
|---|---|---|
| *Serpinh1* | -0.79 | $8.2\times10^{-3}$ |
| *Hspa5* | -0.80 | $1.2\times10^{-2}$ |
| *Mdk* | -0.60 | $1.3\times10^{-2}$ |
| *Ptprk* | 0.57 | $1.6\times10^{-2}$ |
| *1700018A04Rik* | 1.00 | $3.1\times10^{-2}$ |
| *Lcorl* | 0.48 | $3.7\times10^{-2}$ |

**B**

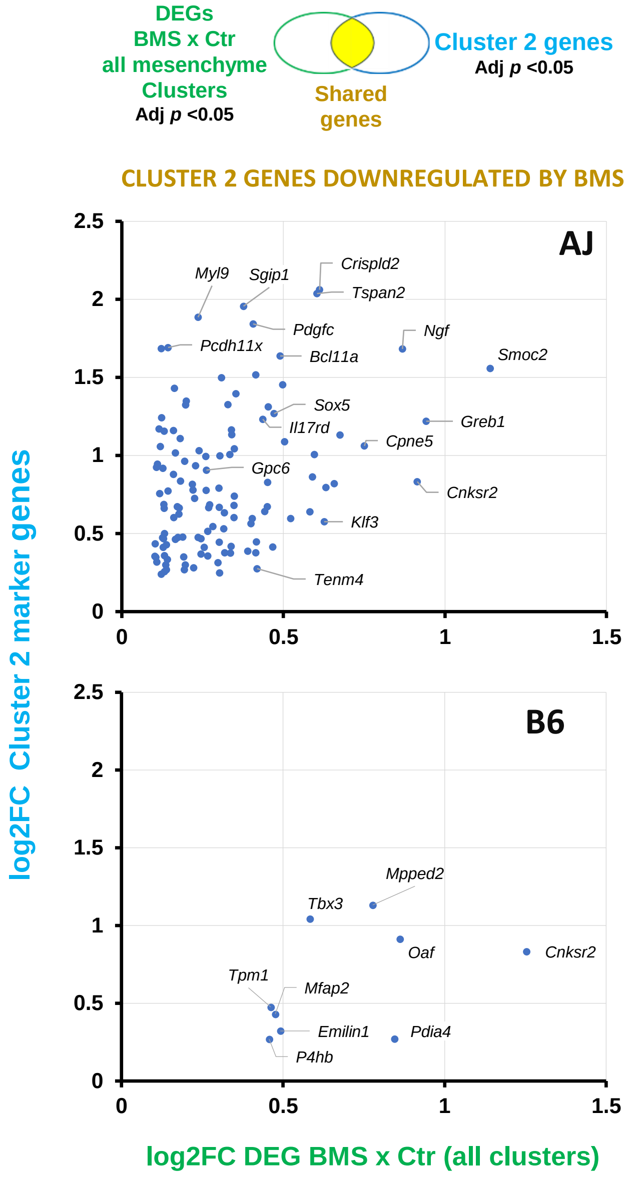


**C**

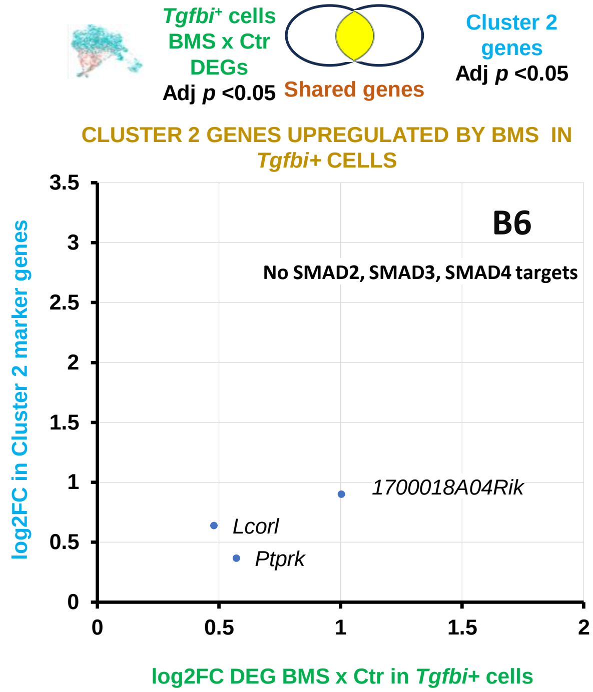


**Suppl. Figure 7**
**Otoshi et al.**
